# RNA polymerase sliding on DNA can couple the transcription of nearby bacterial operons

**DOI:** 10.1101/2023.02.10.528045

**Authors:** Debora Tenenbaum, Koe Inlow, Larry Friedman, Anthony Cai, Jeff Gelles, Jane Kondev

## Abstract

DNA transcription initiates after an RNA polymerase (RNAP) molecule binds to the promoter of a gene. In bacteria, the canonical picture is that RNAP comes from the cytoplasmic pool of freely diffusing RNAP molecules. Recent experiments suggest the possible existence of a separate pool of polymerases, competent for initiation, which freely slide on the DNA after having terminated one round of transcription. Promoter-dependent transcription reinitiation from this pool of post-termination RNAP may lead to coupled initiation at nearby operons, but it is unclear whether this can occur over the distance- and time-scales needed for it to function widely on a bacterial genome in vivo. Here, we mathematically model the hypothesized reinitiation mechanism as a diffusion-to-capture process and compute the distances over which significant inter-operon coupling can occur and the time required. These quantities depend on previously uncharacterized molecular association and dissociation rate constants between DNA, RNAP and the transcription initiation factor *σ*^70^; we measure these rate constants using single-molecule experiments in vitro. Our combined theory/experimental results demonstrate that efficient coupling can occur at physiologically relevant *σ*^70^ concentrations and on timescales appropriate for transcript synthesis. Coupling is efficient over terminator-promoter distances up to ∼ 1, 000 bp, which includes the majority of terminator-promoter nearest neighbor pairs in the *E. coli* genome. The results suggest a generalized mechanism that couples the transcription of nearby operons and breaks the paradigm that each binding of RNAP to DNA can produce at most one messenger RNA.

**SIGNIFICANCE STATEMENT:** After transcribing an operon, a bacterial RNA polymerase can stay bound to DNA, slide along it, and reini-tiate transcription of the same or a different operon. Quantitative single-molecule biophysics experiments combined with mathematical theory demonstrate that this reinitiation process can be quick and efficient over gene spacings typical of a bacterial genome. Reinitiation may provide a mechanism to orchestrate the transcriptional activities of groups of nearby operons.

## INTRODUCTION

The core RNA polymerase (RNAP) in bacteria, which is composed of five subunits (*β, β*′, *α*^*I*^, *α*^*II*^, and *ω*), can catalyze the synthesis of RNA, but cannot recognize specific promoter sequences. To recognize promoters, RNAP must first bind an initiation factor such as the *E. coli* housekeeping *σ*^70^, forming an RNAP holoenzyme (*σ*^70^RNAP) ^1–3^. DNA transcription initiates when *σ*^70^RNAP is recruited from cytoplasm to bind to the promoter region of a gene. Controlling this process is thought to be the principal means through which transcription repressors and activators modulate gene transcription.

Bacterial metabolism requires the coordinated expression of multiple genes^4^. A basic way in which this coordination is achieved in bacterial cells is by the organization of functionally related genes into operons, which are groups of consecutive genes that can be transcribed from the same promoter^5,6^. Functionally related operons often reside in contiguous regions of the bacterial genome^7^. Proximally located operons show higher levels of correlated expression than distant operons in *E. coli*^8–10^. The same is true of closely spaced genes in eukaryotes^11,12^.

While specific groups of bacterial operons may have correlated activities simply because they have common regulatory proteins (e.g., alternative sigma factors), there are also proximity-based mechanisms that can couple transcription of adjacent operons. Terminator readthrough, in which the RNAP fails to read a terminator signal and keeps elongating the mRNA molecule, can generate the joint transcription of codirectional neighboring operons. Transcription-coupled DNA supercoiling^13^ can induce coupled transcription of divergently transcribed genes^14,15^.

Recently, a new mechanism of proximity-based transcription coupling was observed. Using single-molecule microscopy, Harden et al.^16^ and Kang et al.^17,18^ observed that RNAP can remain bound to DNA after termination for at least hundreds of seconds in vitro. This post-termination RNAP-DNA complex may retain a partially open bubble in the DNA^19^. The retained RNAP exhibits one-dimensional diffusive sliding over hundreds or thousands of base pairs along the DNA. In the presence of *σ*^70^ in solution, the sliding RNAP can re-initiate transcription at a nearby promoter. This post-termination behavior of bacterial RNAP may couple transcription of nearby operons in a way that is dependent on both the distance between the two transcription units and the available concentration of *σ*^70^. Genome-wide transcription measurements are consistent with this mechanism, but do not prove that it operates in vivo in both *E. coli* and *B. subtilis*^16^.

In this work, we test whether RNAP post-termination sliding followed by *σ*^70^ rebinding can efficiently couple the transcription of nearby operons. First, we mathematically model the mechanism as a diffusion-to-capture process, in which the association of a *σ*^70^ molecule with the sliding RNAP is required for re-initiation at a nearby promoter sequence. Next, we use single-molecule microscopy experiments under conditions designed to mimic the ionic composition of bacterial cytoplasm to measure the values of the model kinetic parameters. Finally, we input the measured values into the model to predict the distances and times over which post-termination sliding of RNAP could couple expression of neighboring genes.

## RESULTS

### Model of operon coupling by sliding RNAP

We model transcriptional coupling between proximal operons using the sliding RNAP mechanism depicted in Fig. 1. The distance between the terminator of the first operon and the promoter of the second operon is *d*. Upon reaching the terminator sequence T of the primary operon (at time *t* = 0), the RNAP releases an RNA transcript but remains non-specifically bound, enabling it to diffuse along the DNA with a diffusion coefficient *D*. During this time interval of sliding, the RNAP can either dissociate from the DNA with rate *k*_off_, or bind a *σ*^70^ molecule from solution with a rate *k*_b_[*σ*^70^], where [*σ*^70^] denotes the free *σ*^70^ solution concentration. In the latter case, the RNAP-*σ*^70^ complex continues diffusing along the DNA molecule, and can dissociate with a rate *k*_off,s_, or can encounter and be captured by the promoter for the secondary operon. We define the time it takes the captured RNAP-*σ*^70^ complexes to find the secondary promoter as the search time *t*_f_. In this mechanism, *σ*^70^ can have conflicting effects because it can stimulate RNAP dissociation from DNA via the *k*_off,s_ step and yet is also required for secondary promoter capture.

**Figure 1:**
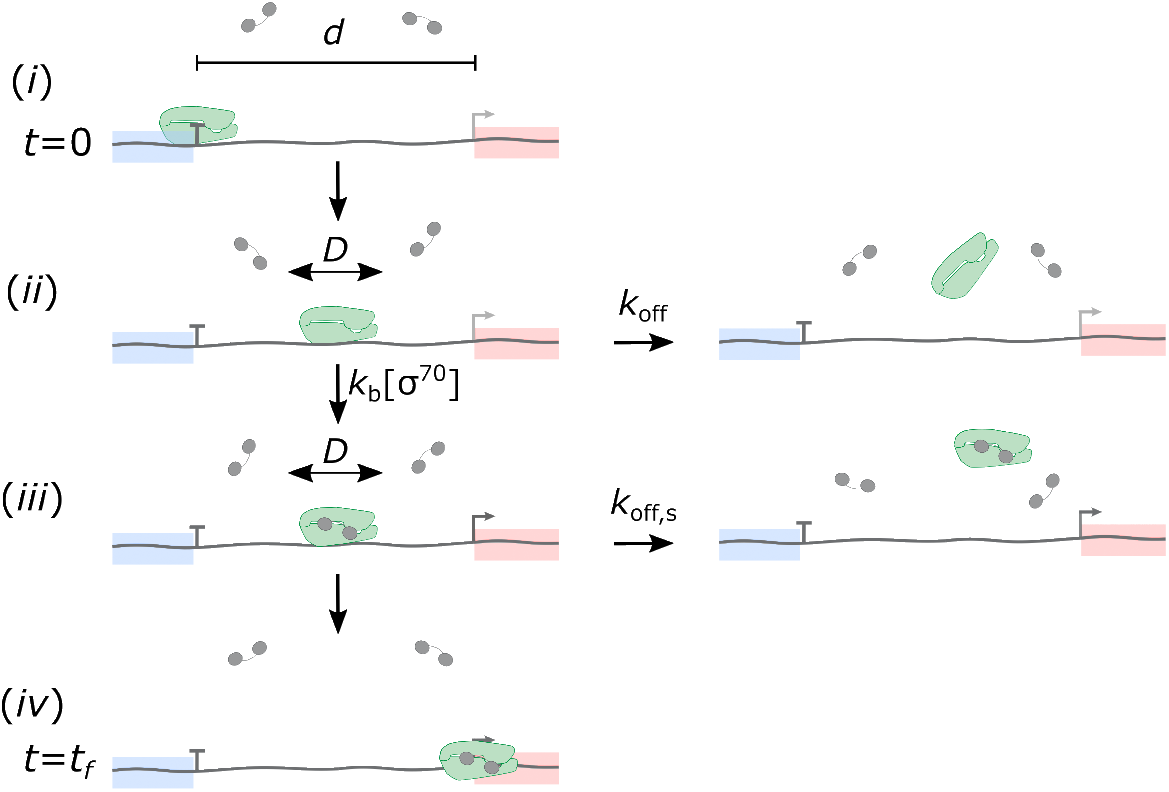
Model of operon coupling by sliding RNAP. (i) An RNAP molecule (green) terminates transcription of the primary operon (blue), and (ii) starts sliding along the DNA molecule with a diffusion constant *D*. (iii) While sliding, the RNAP can either dissociate from the DNA with rate *k*_off_, or bind *σ*^70^ (gray) with rate *k*_b_[*σ*^70^]. (iv) After binding of *σ*^70^, the RNAP-*σ*^70^ complex can either dissociate with a rate *k*_off,s_, or find the promoter (bent arrow) for the secondary operon (pink), which is located is at a distance *d* along the DNA from the primary operon terminator (T).

For simplicity, we assume that the binding of *σ*^70^ to sliding RNAP is irreversible. This is equivalent to assuming that the unbinding of *σ*^70^ from the sliding RNAP is significantly slower than the dissociation of the RNAP-*σ*^70^ complex from the DNA. This assumption is reasonable, given previous measurements of *σ*^70^ dissociation from free^20,21^ and DNA-bound RNAP^17^. Also for simplicity, we assume that the diffusion coefficient on DNA of the RNAP-*σ*^70^ complex and RNAP are the same.

In this work we will refer to the complex that is formed by binding of *σ*^70^ to DNA-bound RNAP as RNAP-*σ*^70^-DNA and to the complex formed by binding *σ*^70^RNAP holoenzyme to DNA as holoenzyme-DNA. It is not currently known whether these different orders of assembly produce complexes with the same structure (see Discussion).

### Calculation of coupling efficiency

To quantify how transcriptional coupling by sliding RNAP changes with varying distance between operons and with [*σ*^70^], we define the coupling efficiency (*E*) as the probability that an RNAP molecule, which terminates transcription of the primary operon, reaches the promoter of the secondary promoter by the sliding RNAP mechanism. To reach the secondary promoter, (i) the sliding RNAP has to bind a *σ*^70^ molecule from solution before falling off the DNA, and (ii) the RNAP-*σ*^70^ complex then has to reach the secondary promoter before falling off the DNA. The probability of (i), *P*_bind_, is given by the partition ratio

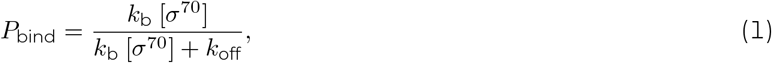

while the probability of (ii), *P*_find_, can be computed as the probability that a RNAP-*σ*^70^ complex remains bound to DNA for at least the time needed to find the secondary promoter before dissociating.

To determine *P*_find_, we combine the distribution of times it takes RNAP-*σ*^70^ complexes to encounter the secondary promoter for the first time, *p*_first passage_(*t*), and the probability that the complex will stay bound on the DNA long enough for the encounter to happen. Thus, *P*_find_ is given by

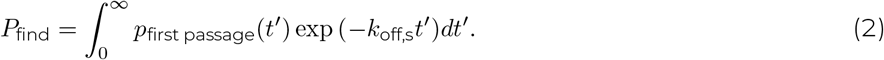

Here we use uppercase *P* to refer to probabilities, and lowercase *p* to represent probability density functions (PDFs).

RNAP terminates transcription of the primary operon at *x* = 0 and starts performing a one-dimensional random walk along the DNA with diffusion coefficient *D*. How long it takes for a RNAP-*σ*^70^ complex to first encounter the secondary promoter will depend on its position on the DNA (*x*_b_) when it starts the search, i.e., when *σ*^70^ binds the sliding RNAP molecule. *x*_b_ in turn depends on how long after termination at the primary terminator *σ*^70^ binds (*t*_b_). At a time *t*_b_ drawn at random from the exponential distribution

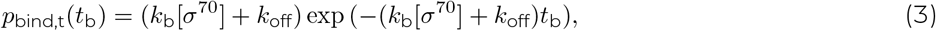

the sliding RNAP will either bind a *σ*^70^ molecule or dissociate from the DNA. If it binds a *σ*^70^ molecule, the binding position will be a random value drawn from

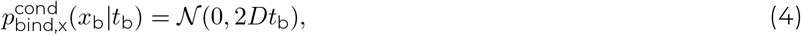

where 𝒩(*μ, var*) describes a normal distribution with mean *μ* and variance *var*. Eq. 4 represents the conditional probability distribution for *x*_b_ given the binding time *t*_b_. Now we can calculate the unconditional distribution of binding positions

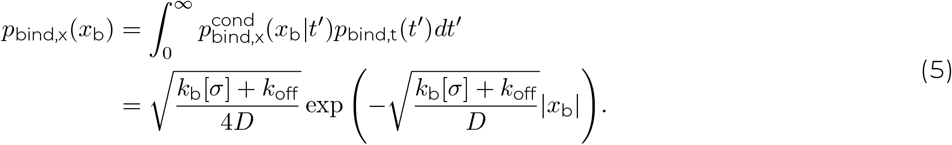

The distribution of times *t*_fp_ it would take the RNAP-*σ*^70^ complex to reach the secondary promoter at *x* = *d* for the first time is then given by the first-passage-time density for a one-dimensional random walk^22^

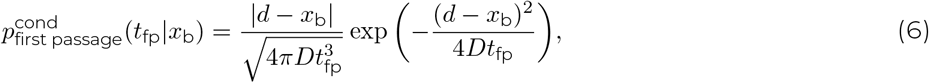

which is conditional on *x*_b_. The unconditional distribution of first passage times is then calculated as

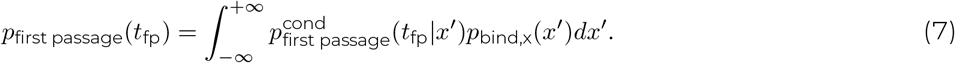

Finally, substituting Eq. 7 into Eq. 2 to get *P*_find_, and multiplying by *P*_bind_ (Eq. 1), we get the expressions in Eq. 8.

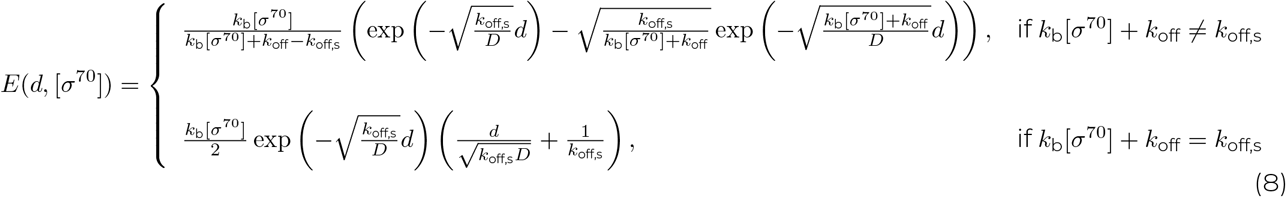

### Coupling regimes and calculation of coupling distance

We can distinguish three different coupling regimes depending on the availability of free *σ*^70^ molecules. For this, we define a critical *σ*^70^ concentration at which the diffusion time intervals available before and after *σ*^70^ binding to RNAP are equal, 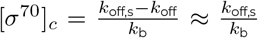. Here we used *k*_off_ << *k*_off,s_ based on prior studies^23^ and our experimental results (see below).

1. When [*σ*^70^] ≫ [*σ*^70^]_*c*_, the coupling efficiency at small *d* is given by *P*_bind_ (Eq. 1) since any RNAP that binds *σ*^70^ will subsequently encounter the promoter. For larger *d* the efficiency decays exponentially,

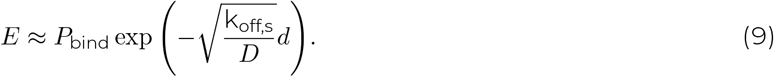

In this regime, we can define the *coupling distance* as the characteristic decay distance of the coupling, 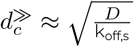.
2. When [*σ*^70^] ≪ [*σ*^70^]_*c*_, the decay is also exponential but in this case,

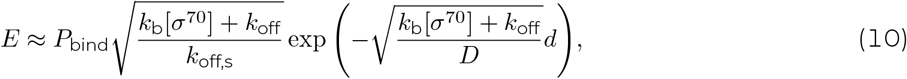

and the characteristic distance is 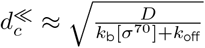. However, in this case 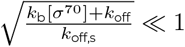, which means that there is no significant coupling between adjacent transcription units no matter the distance between them.
3. For [*σ*^70^] ≈ [*σ*^70^]_*c*_, we get the bottom expression in Eq. 8. Even though it is not exponential, we can still define a coupling distance 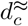 over which the coupling efficiency decays by a factor of *e*. Following the calculations in Appendix S1, we get

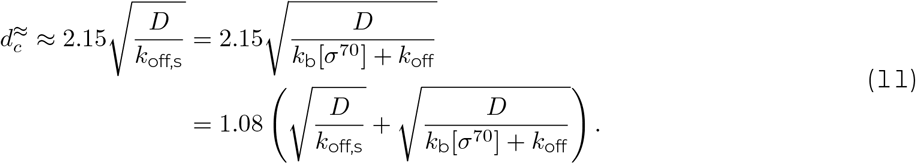

In simple terms, the regimes differ by whether most of the diffusional search for the secondary promoter takes place after (regime 1) or before (regime 2) the binding of *σ*^70^. In addition, given that 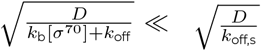 in regime 1, and 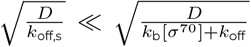 in regime 2, for all three regimes we can then approximate the coupling distance as 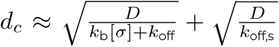, which is roughly the sum of the root mean squared displacements of the RNAP before and after binding a *σ*^70^ molecule (Fig. S1). Efficiency curves as a function of distance scaled by the critical distance are shown for all three regimes in Fig. 2. As expected, when the concentration of *σ*^70^ is well below its critical concentration, the efficiency is small for any distance between the two promoters, while the efficiency can be of order one when [*σ*^70^] is well above [*σ*^70^]_*c*_.

**Figure 2:**
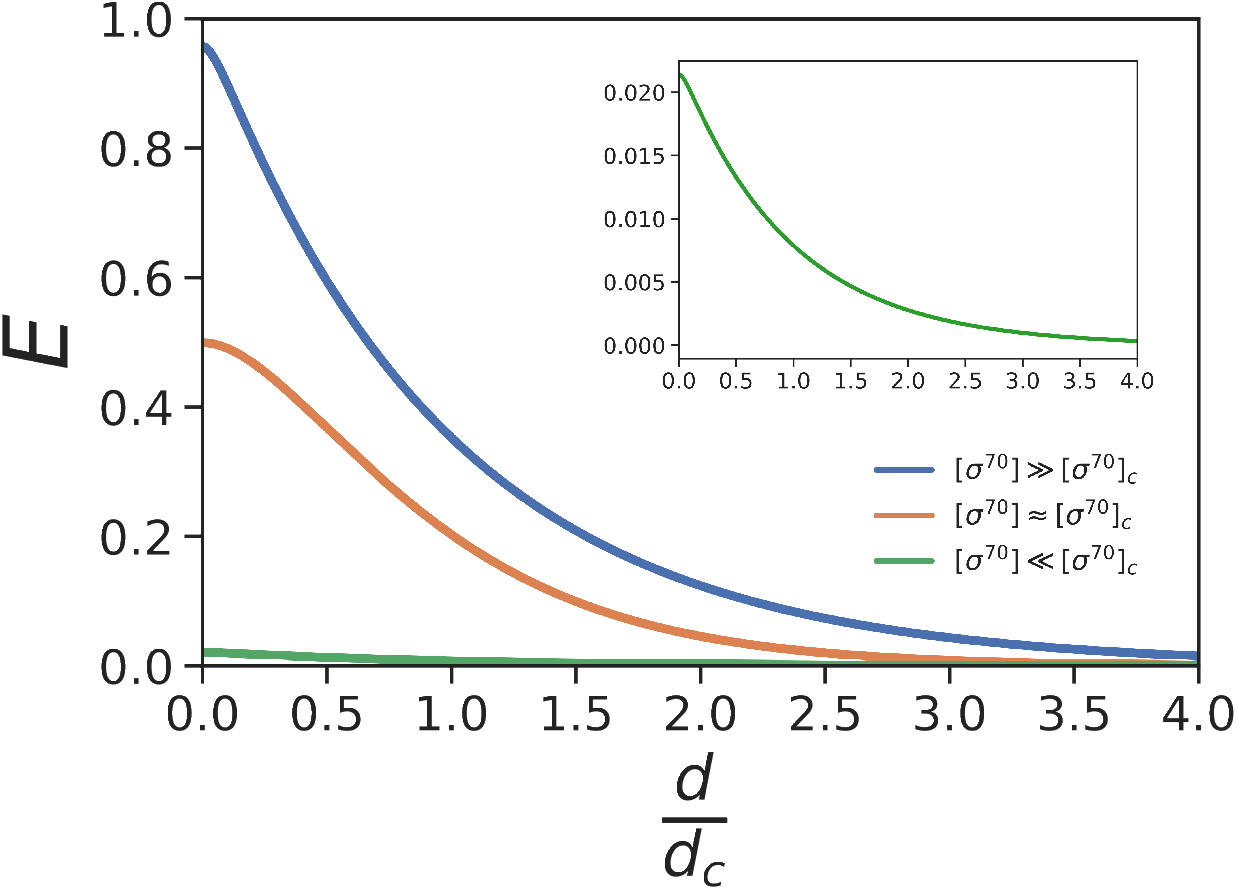
Predicted relationship of Coupling efficiency *E* to the distance between operons. The coupling efficiency is the probability of an RNAP that terminated transcription at the end of the primary operon reaching the secondary promoter. Efficiency curves are shown for the three regimes of *σ*^70^ concentration described in the text, for a chosen set of parameter values: *k*_b_ = 10^7^ M^−1^s^−1^, *k*_off_ = 10^−3^ s^−1^, *k*_off,s_ = 1 s^−1^, *D* = 4 *×*10^4^ bp^2^s^−1^. Curves were calculated using Eq. 8. Distance is represented in units of the coupling distance 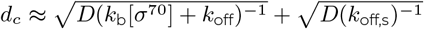. Inset: enlarged plot of the curve for [*σ*^70^] ≪ [*σ*^70^]_*c*_.

### Single-molecule microscopy experiments to measure model parameters

To estimate the coupling efficiency *E*, coupling distance *d*_c_, and search times *t*_f_ that can be achieved via the proposed mechanism (Fig. 1), we need values for the model parameters *D, k*_b_, *k*_off_, and *k*_off,s_.

The diffusion coefficient of RNAP post termination, *D* = (3.9 ± 0.5) × 10^4^ bp^2^s^−1^, was experimentally measured in ref.^16^. Those investigators also sometimes observed a non-diffusing post termination RNAP-DNA complex in their experiments, but attributed this to RNAP binding to the ends of the linear DNA molecules they used.

We performed single-molecule experiments to measure *k*_b_, *k*_off_ and *k*_off,s_. Specifically, we quantified the dwell times of RNAP on promoterless DNA templates in the presence of different concentrations of *σ*^70^ (Fig. 3A). These experiments allow us to measure all three rate constants. This is because at low *σ*^70^ concentrations the measured dwell times are limited by the rate of RNAP dissociation from DNA, at intermediate *σ*^70^ concentrations they are limited by the rate of *σ*^70^ binding to the RNAP-DNA complex, and at high *σ*^70^ concentrations they are limited by the rate of RNAP-*σ*^70^ complex dissociation from DNA.

**Figure 3:**
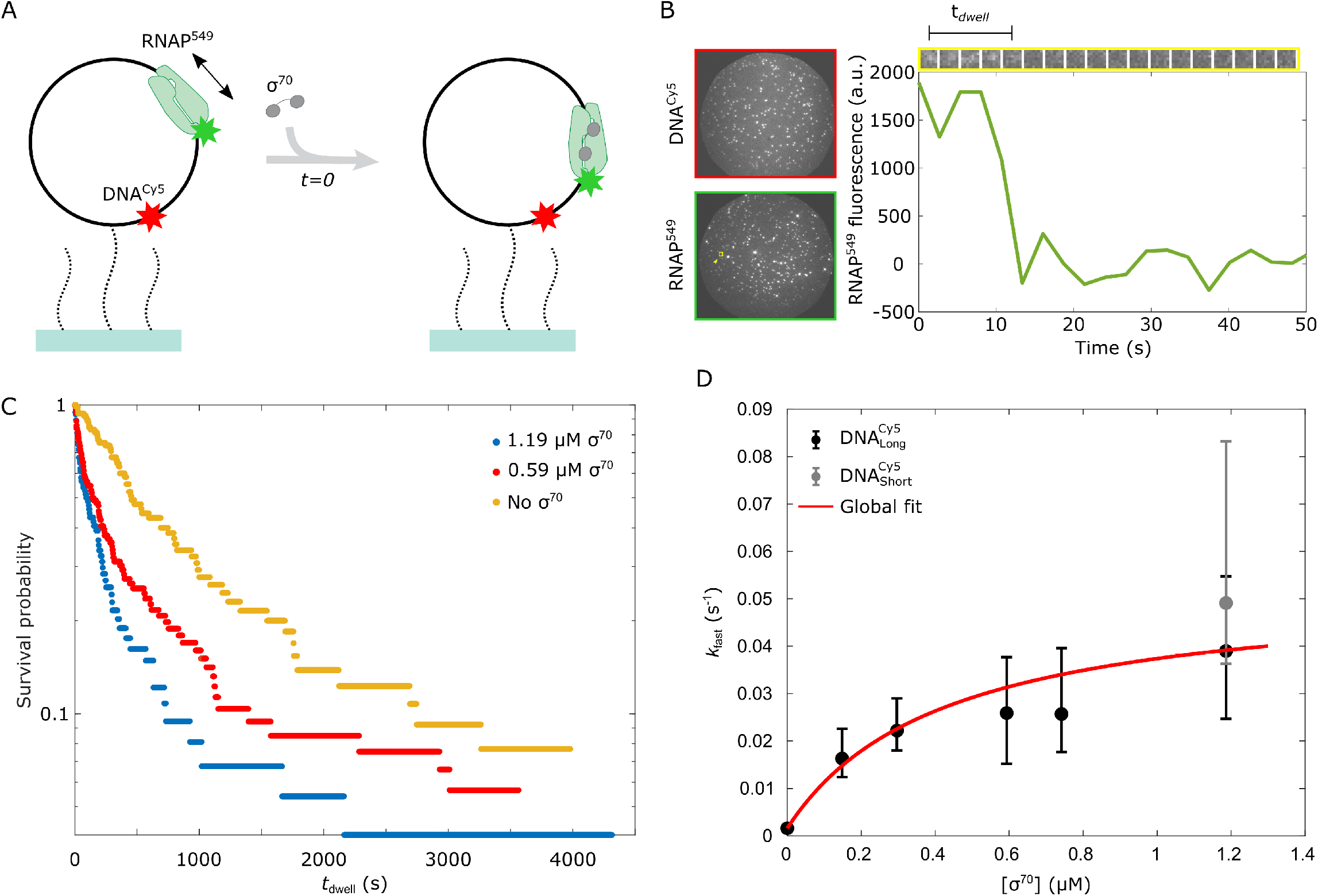
Single-molecule experiments to measure *k*_b_, *k*_off_, and *k*_off,s_. **a.** Experiment schematic. Fluorescently labeled, promoterless circular DNA templates (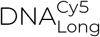; black circles) were tethered to the surface of a glass flow chamber (blue) through polyethylene glycol linkers (dotted black curves). The chamber was then incubated with fluorescently labeled core RNAP (RNAP^549^) to form RNAP-DNA nonspecific complexes, which correspond to the post-termination complex. In each of six experiments, a different concentration of *σ*^70^ (gray) was introduced at *t* = 0, and the lifetime of each RNAP^549^ that colocalized with a surface-tethered DNA molecule was monitored by single-molecule fluorescence microscopy. **B**. Example of experiment record. Left: 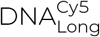 and RNAP^549^ fluorescence images of the same field of view (65 *μ*m diameter) upon introducing 1.19 *μ*M *σ*^70^ at *t* = 0. Right: Time record excerpt of RNAP^549^ fluorescence at the location of a single DNA molecule. Gallery shows 5 *×*5 pixel images centered on the DNA molecule; graph shows the summed, background-corrected intensity of the 3*×* 3 pixels centered on the DNA. *t*_dwell_ represents the duration of the fluorescent spot. **C**. *t*_dwell_ survival probability distributions in the presence of 0, 0.59, and 1.19 *μ*M *σ*^70^. **D**. Rates (with 68% C.I.s) of *σ*^70^-dependent dissociation of RNAP^549^ from 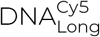 (black) and 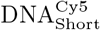 (gray) as a function of *σ*^70^ concentration, and global fit (red; see eq. (12) and accompanying text).

For experimental convenience, we did not use core RNAP-DNA complexes that were formed after termination of transcription. Instead, we directly formed sequence non-specific RNAP-DNA complexes by adding core RNAP to a DNA that lacks known promoter sequences. The two types of complexes have the same protein composition and have similar properties: both are long-lived, in both RNAP slides on DNA, both are sensitive to the polyanion heparin^16^, and both are rapidly disassembled by the bacterial SNF2 ATPase RapA^24^.

To implement these experiments we designed and synthesized a biotinylated, 3,033 bp circular DNA lacking known promoter sequences that was labeled with the red-excited dye Cy5 (we refer to this construct as 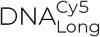). Circular DNAs were used to avoid possible binding of RNAP to DNA ends^25^, which are largely non-physiological since the *E. coli* chromosome is circular.

We immobilized 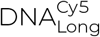 molecules on the surface of a glass flow chamber via a biotin-streptavidin linkage (Fig. 3A). We then incubated the chamber with a solution containing *E. coli* core RNAP labeled with a green-excited dye (RNAP^549^) for ∼ 10 min, and washed it out at time *t* = 0 with a solution containing *σ*^70^ in the 0 to 1.2 *μ*M range. Single-molecule total internal reflection microscopy was performed with alternating red and green excitation for observation of 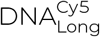 and RNAP^549^, respectively. An example of the fluorescence records used for extracting the dwell times of the RNAP^549^ molecules on the 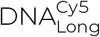 template for each experiment is shown in Fig. 3B.

Given that these experiments study sequence-nonspecific interactions between RNAP^549^ and 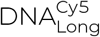, it was expected that multiple RNAP^549^ molecules could be bound to the same 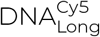 template simultaneously. To reduce complications in the dwell time measurements arising from multiple RNAP^549^ molecules bound to the same template, we restricted the analysis to only those 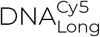 locations with a single colocalized RNAP^549^. The number of RNAP^549^ molecules bound to each 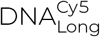 template was quantified by counting the number of decreasing steps present in the RNAP^549^ fluorescence intensity records (Fig. S2 and Appendix S3).

### Distributions of RNAP dwell times on DNA

Example dwell time probability distributions of RNAP^549^ on promoterless circular DNA templates for different concentrations of *σ*^70^ are shown in Fig. 3C. Consistent with the results in^23^, *σ*^70^ accelerates the dissociation of RNAP from DNA.

In the absence of promoter sequences in the DNA, the model in Fig. 1 predicts that the dwell time distributions for RNAP obtained in the limits of low and high [*σ*^70^] are exponential. Theoretically, at intermediate [*σ*^70^] the dwell time distributions are non-exponential, due to the presence of two sequential steps (*k*_b_ and *k*_off,s_). Still, for reasonable values of the rate constants the distribution is well approximated by an exponential and the effective rate constant has a hyperbolic dependence on [*σ*^70^],

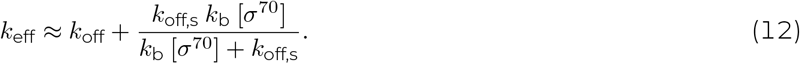

However, the experimental distributions are in fact described not by a single exponential, but by the sum of multiple exponential components with very different rate constants. Specifically, for the experiments in which [*σ*^70^] *>* 0 the distributions are well fit by a sum of three exponentials with characteristic rates *k*_slow_, *k*_inter_, and *k*_fast_ (Fig. S3). This suggests that in these experiments there are at least three types of RNAP-*σ*^70^-DNA complexes. The resulting fits, obtained using a maximum likelihood method, are shown in Fig. S4, and the fit parameters are summarized in Table 1. In all cases, non-specific binding of RNAP^549^ to the chamber surface was minimal (Table S1), and therefore was not considered when fitting the data.

**Table 1:** Parameters for fits to dwell time distributions of RNAP^549^-promoterless DNA complexes at different *σ*^70^ concentrations.

| $[\sigma^{70}]$ ($\mu\text{M}$) | N | N<sub>d</sub> | $a_{\text{fast}} (\times 10^{-1})$ | $a_{\text{inter}} (\times 10^{-1})$ | $k_{\text{fast}} (\times 10^{-2} \text{ s}^{-1})$ | $k_{\text{inter}} (\times 10^{-3} \text{ s}^{-1})$ | $k_{\text{slow}} (\times 10^{-4} \text{ s}^{-1})$ | DNA preparation |
| --- | --- | --- | --- | --- | --- | --- | --- | --- |
| 0 | 65 | 60 | 8.6(6.8 – 9.4) | - | 0.16(0.13 – 0.22) | - | 1.59(0 – 3.99) | DNA<sup>Cy5</sup><sub>Long</sub>, preparation 1 |
| 0.15 | 60 | 42 | 1.2(0.4 – 2.0) | 1.6(0.4 – 2.9) | 1.63(1.24 – 2.26) | 3.11(2.27 – 4.52) | 1.90(1.44 – 2.33) | DNA<sup>Cy5</sup><sub>Long</sub>, preparation 2 |
| 0.30 | 139 | 85 | 1.8(1.3 – 2.1) | 1.3(0.7 – 3.8) | 2.22(1.80 – 2.90) | 1.23(0.53 – 2.04) | 1.27(0.48 – 1.52) | DNA<sup>Cy5</sup><sub>Long</sub>, preparation 2 |
| 0.59 | 106 | 100 | 3.4(2.5 – 4.8) | 5.4(4.1 – 6.2) | 2.59(1.52 – 3.77) | 2.61(1.87 – 3.37) | 2.18(0.26 – 3.63) | DNA<sup>Cy5</sup><sub>Long</sub>, preparation 1 |
| 0.74 | 106 | 60 | 1.8(1.2 – 2.2) | 2.0(1.0 – 5.2) | 2.57(1.77 – 3.96) | 0.81(0.34 – 1.72) | 1.05(0 – 1.54) | DNA<sup>Cy5</sup><sub>Long</sub>, preparation 3 |
| 1.19 | 74 | 71 | 3.4(2.3 – 4.9) | 5.8(4.4 – 6.8) | 3.90(2.47 – 5.47) | 3.88(2.45 – 5.26) | 1.61(0 – 3.24) | DNA<sup>Cy5</sup><sub>Long</sub>, preparation 1 |
| 1.19 | 76 | 68 | 5.8(4.0 – 6.7) | 3.0(2.0 – 4.7) | 4.91(3.63 – 8.32) | 5.77(4.03 – 10.09) | 0.72(0 – 1.56) | DNA<sup>Cy5</sup><sub>Short</sub> |

The models used for the fit are described in Methods (Eqs. 13 and 14). N is the number of DNA sites with co-localized RNAP that were used in the analysis. N<sub>d</sub> is the number of DNA sites for which the co-localized RNAPs disappeared before the end of the experiment. The values are presented with 68% CI. In some cases, the lower confidence limit on $k_{\text{slow}}$ is poorly defined because $k_{\text{slow}}^{-1}$ exceeds the duration of the experiment. DNA<sup>Cy5</sup><sub>Long</sub> preparations 1, 2, and 3 were made by the same method on different occasions.

For the experiments with [*σ*^70^] *>* 0, *k*_fast_ showed a hyperbolic dependence of [*σ*^70^] (Fig. 3D); *k*_inter_ and *k*_slow_ did not (Table 1). Therefore, we hypothesize that the fastest component corresponds to *σ*^70^-induced dissociation of RNAP^549^ bound to DNA, *k*_eff_ = *k*_fast_. This suggests that at low *σ*^70^ concentrations binding of *σ*^70^ to the sliding RNAP is rate-limiting so that the dissociation rate of RNAP from DNA increases linearly with *σ*^70^ concentration, while at high concentrations the dissociation of the RNAP-*σ*^70^ complex from DNA becomes limiting, and the dissociation rate saturates. Possible origins of the longer-lived RNAP-DNA complexes with [*σ*^70^]-independent dissociation rates *k*_inter_ and *k*_slow_ are discussed in the Appendix S4.

To confirm that *k*_fast_ depends on [*σ*^70^] and not on the DNA template used, we repeated the 1.19 *μ*M *σ*^70^ experiment using a different DNA template, the promoterless 586 bp circular 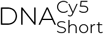. Similar values were obtained for *k*_fast_ for both templates (Table 1, Fig. 3D), supporting the idea that *k*_fast_ depends on *σ*^70^ concentration, and not on the length or sequence of the DNA template used.

Two characteristic rates were observed for the experiment with [*σ*^70^] = 0. The slower one is similar to the values of *k*_slow_ observed in the presence of *σ*^70^ (Table 1). The faster one is similar to the mean dissociation rate observed for the post-termination RNAP-DNA complex in the absence of *σ*^70^, as well as to the mean dissociation rate observed for core RNAP sequence-nonspecifically bound to DNA^16^. Therefore, we assume that the faster rate corresponds to the dissociation rate of RNAP^549^ from DNA in the limit where [*σ*^70^] = 0.

### Extraction of model parameters *k*_b_, *k*_off_, and *k*_off,s_

Having established the hyperbolic dependence of the *σ*^70^-induced dissociation rate *k*_fast_ on *σ*^70^ concentration, we can now determine the values for the Fig. 1 model parameters *k*_b_, *k*_off_, and *k*_off,s_. For this, we jointly fit the data from experiments at different *σ*^70^ concentrations to a global model that incorporates our conclusions about the origins of the different components of the dwell time distributions (see Appendix S4). The *σ*^70^-independent rates *k*_inter_ and *k*_slow_ were globally fit for all six experiments performed with 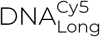 (Table S3 and Fig. S8). A separate set of parameters 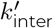 and 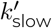 were obtained by fitting the dwell time distribution from the experiment with 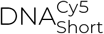. The global model explicitly included the [*σ*^70^] dependency of *k*_fast_ (Eq. 12, where *k*_eff_ = *k*_fast_).

The model fit well to the data (Fig. S8) and gave well-constrained values for the rate constants (Table 2). The rate constants, together with the diffusion coefficient *D* measured in ref.^16^ provide the information needed to calculate the extent and kinetics of operon coupling.

**Table 2:** Global model parameters.

| Parameter | Description | Value (68% C.I.) | Source |
| --- | --- | --- | --- |
| $D$ | Diffusion coefficient of sliding RNAP | $3.9 \times 10^4 \text{ bp}^2\text{s}^{-1}\dagger$ | 16 |
| $k_b$ | Binding rate of $\sigma^{70}$ to sliding RNAP | $1.2(0.7 - 2.7) \times 10^5 \text{ M}^{-1}\text{s}^{-1}$ | Fit |
| $k_{\text{off}}$ | Dissociation rate of RNAP before binding $\sigma^{70}$ | $1.6(1.3 - 1.9) \times 10^{-3} \text{ s}^{-1}$ | Fit |
| $k_{\text{off},s}$ | Dissociation rate of RNAP-$\sigma^{70}$ complex | $5.1(3.4 - 12.2) \times 10^{-2} \text{ s}^{-1}$ | Fit |

† Kang et al<sup>17</sup> measured a somewhat lower value corresponding to $0.8 \times 10^4 \text{ bp}^2\text{s}^{-1}$ at similar ionic strength but in a different buffer.

### Extent of operon coupling by sliding RNAP

Operon coupling cannot be biologically functional if it takes an infeasibly long time for RNAP to find the secondary promoter after terminating transcription at the terminator of the primary operon. To compute the distribution of search times, we used the experimental results for *D, k*_b_, *k*_off_, and *k*_off,s_ to simulate the mechanism for a realistic *σ*^70^ concentration and terminator-promoter spacing (Fig. 4A). The distribution of the search times is roughly exponential with a mean ⟨*t*_f_⟩ ∼ 7 s, which is comparable to the time for transcription initiation at well-studied promoters (a few seconds to a few minutes^26,27^). This indicates that transcription re-initiation by sliding RNAP is capable of effectively increasing expression of the secondary operon.

**Figure 4:**
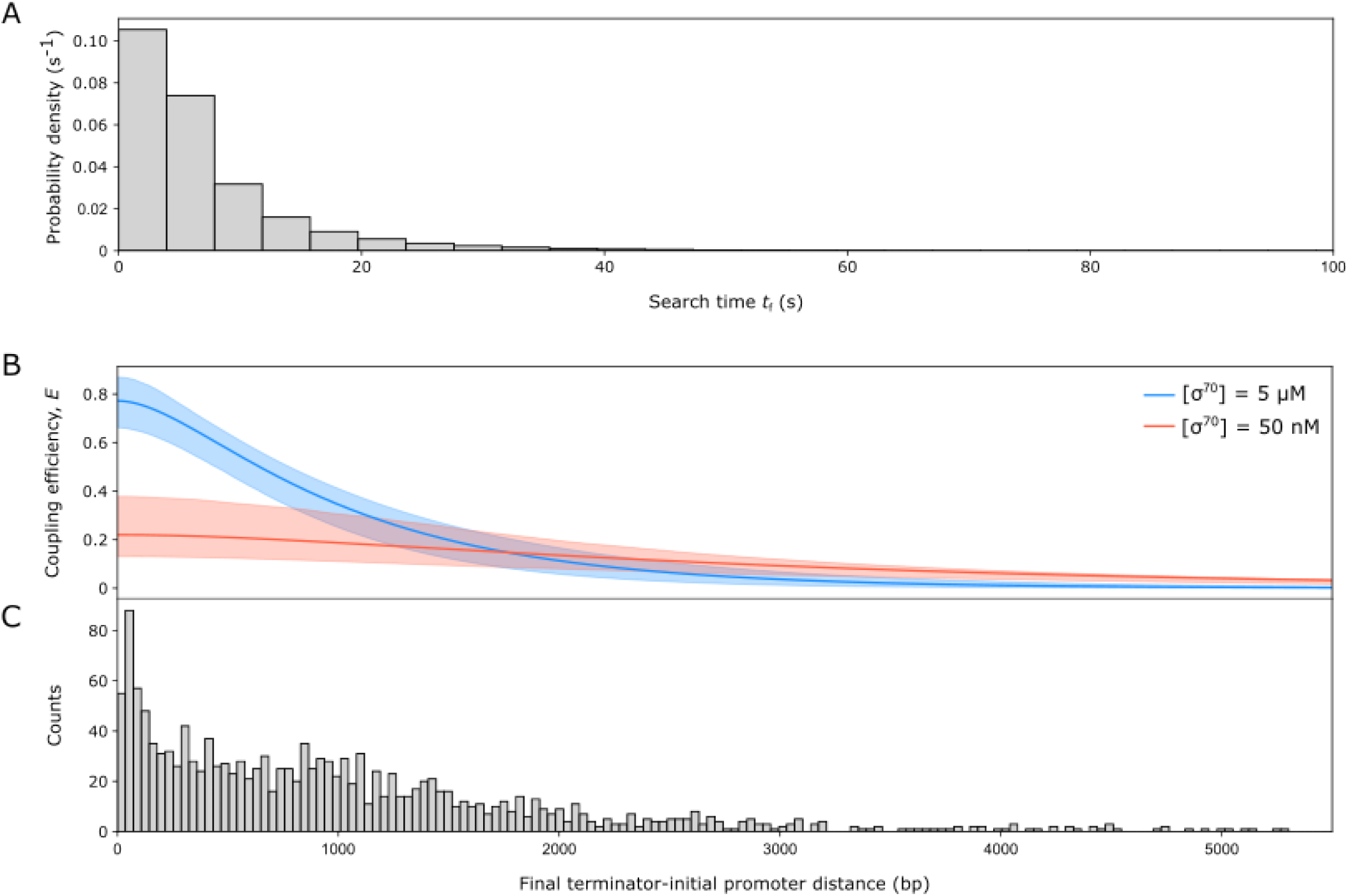
Extent of operon coupling predicted by the sliding RNAP model, using the kinetic parameter values from Table 2. **a**. Distribution of search times that end in a promoter encounter *t*_f_ obtained by simulating the model in Figure 1 for *d* = 600 bp and [*σ*^70^] = 5 *μ*M. **b**. Coupling efficiency dependence on the distance *d* between primary operon terminator and secondary operon promoter, for two possible free *σ*^70^ concentrations. Shaded areas show the 68% C.I.s. **c**. Distribution of distances between operon final terminators and the nearest operon initial promoter in the *E. coli* genome determined from data in ref.^29^.

Inputting the experimental results for *D, k*_b_, *k*_off_, and *k*_off,s_ into Eq. 8, we can predict the value of the efficiency as a function of the distance between operons and the *σ*^70^ concentration. The total concentration of *σ*^70^ in *E. coli* is on order 10 *μ*M^28^, but its availability is highly regulated through sequestration by anti-*σ* factors, whose activity is also tightly regulated. This means that at any time, the free *σ*^70^ concentration could be anything below roughly 10 *μ*M. Thus, the free *σ*^70^ concentration in the cell could be either above or below [*σ*^70^]_*c*_, which we calculate to be 0.4 *μ*M. To test whether the model predicts appreciable coupling at typical operon spacings, we calculated the predicted coupling efficiency as a function of the distance *d* between the primary terminator and the secondary promoter at high and low *σ*^70^ concentrations (Figure 4B). At 5 *μ*M *σ*^70^, this calculation predicts efficient coupling at distances *d* up to 1, 000 bp. At a much lower free *σ*^70^ concentration of 50 *n*M, the model predicts a smaller but still significant amount of coupling on this distance scale, with less dependence on operon spacing. Regulation of the free *σ*^70^ concentration would therefore allow the cell to vary the amount of coupling in response to internal and environmental conditions.

The spacing between the final terminator of an operon and the nearest operon initial promoter has a broad distribution in the *E. coli* genome (Figure 4C). Nevertheless, based on our calculations, a large fraction of these pairs are capable of efficient coupling by sliding RNAP. For example, 52% of the terminator-promoter pairs are at distances where the coupling efficiency is at least 50% at [*σ*^70^] = 5 *μ*M. In other words, at this *σ*^70^ concentration (and in general when [*σ*^70^] *>>* [*σ*^70^]_*c*_ ≈ 0.4 *μ*M) the predicted critical distance *d*_*c*_ ≈ 1, 000 bp is of the same order of magnitude as the typical inter-operon distance (median 600 bp). This could allow many pairs of adjacent operons in the genome to be coupled, while at the same time enabling other operons to be transcribed independently of their neighbors, depending on the terminator-promoter spacing. Thus, the model predicts significant coupling under relevant cellular conditions and predicts that coupling can be regulated by tuning these conditions.

The model makes the simplifying assumption that every encounter of the RNAP-*σ*^70^ complex with a promoter is productive and leads to synthesis of a transcript. To the extent that not all encounters are productive, the model will overestimate efficiency (Fig. 4B) and underestimate search time (Fig. 4A).

## DISCUSSION

Using a combination of theory, stochastic simulations, and single-molecule microscopy experiments, we characterized the potential spatial and temporal reach of transcriptional coupling between adjacent operons mediated by diffusive sliding of RNAP that remains bound to DNA following transcription termination. We predict that *σ*^70^ has both stimulatory and inhibitory effects on reinitiation. The stimulatory effect arises from the fact that only RNAP with bound *σ*^70^ can recognize the secondary promoter. On the other hand, we show that RNAP-*σ*^70^ has only a short lifetime on DNA, during which the sliding *σ*^70^RNAP must find a promoter on the fly to reinitiate transcription. Despite the latter difficulty, we show that reinitiation is expected to be common and efficient for physiological ranges of terminator-promoter spacings and *σ*^70^ concentrations. Thus, our results show that the proposed reinitiation mechanism is consistent with experiments that demonstrate reinitiation in vitro^16,17^ and in vivo^16^.

To quantitatively define the reinitiation process, we measured three previously uncharacterized rate constants: the second-order rate constant for binding of *σ*^70^ to the RNAP-DNA complex, *k*_b_ = 1.2 × 10^5^ M^−1^s^−1^, the rate constant for the dissociation of the RNAP-DNA complex, *k*_off_ = 1.6 × 10^−3^ s^−1^, and the rate constant for the dissociation of the RNAP-*σ*^70^-DNA complex, *k*_off,s_ = 5.1 × 10^−2^ s^−1^. The value obtained for *k*_b_ is an order of magnitude smaller than the rate constant of formation of a stable *σ*^70^-RNAP complex in the absence of DNA, 1.5 × 10^6^ M^−1^s^−1 21^, which suggests that the presence of bound DNA significantly impedes *σ*^70^ association with RNAP. The value obtained for *k*_off,s_ is an order of magnitude smaller than the value obtained for 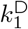, the dissociation rate of the RNAP holoenzyme-DNA complex (see Appendix S4 and Table S2). This suggests that the RNAP-*σ*^70^-DNA complex (formed by RNAP-DNA binding *σ*^70^ from solution) and the holoenzyme-DNA complex (formed by mixing *σ*^70^RNAP with non-promoter DNA) have different conformations, despite them having the same protein and DNA constituents. It is possible that in the two complexes different subsets of *σ*^70^ subregions interact with RNAP and/or DNA. More information, kinetic and structural, will be required to understand these differences.

The search for target sequences by proteins sliding on DNA has been demonstrated both in vitro and in vivo (e.g.,^30,31^). Post-termination sliding of core RNAP on DNA is atypically slow compared to a sample of other DNA binding proteins^16,32^, possibly because RNAP maintains an open bubble of non-base-paired DNA in the post-termination RNAP-DNA complex^19^. The presence in cells of sliding RNAP molecules that may take on order of 10 s after termination to reinitiate transcription (Fig. 4A) is consistent with demonstration of a substantial population in vivo of slowly diffusing RNAP molecules that are neither bound to a fixed site on DNA nor freely diffusing in solution^33,34^.

Rapid, efficient reinitiation of transcription through sliding of post-termination RNAP over relevant genomic distances may have significant implications for transcription homeostasis and regulation in both natural and engineered genomes. Under particular growth conditions, transcription activity is often concentrated in clusters of genes or operons in confined genomic regions^4,6,7,35,36^. Sliding-mediated reinitiation may help to maintain a localized pool of RNAP molecules that are efficiently reused in these transcriptionally active regions. Indeed, the efficiency of reinitiation by sliding core RNAP compared to conventional initiation by RNAP holoenzyme from solution may be one of the factors that confers a selective advantage to the clustering of functionally related operons. In the context of synthetic biology, reinitiation by sliding might cause problems by giving rise to non-intended connectivity between transcription units that are intended to act modularly, but conversely could be a used as a tool to introduce correlations in designed genetic circuits.

## MATERIALS AND METHODS

### Plasmids

Plasmid pDT4 is identical to pCDW116^16^ except for mutation of CTGGAGTGCG to CTGGAGACCG to introduce a second BsaI site.

### DNA templates

Circular DNA templates 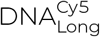 and 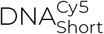 were built by Golden Gate Assembly^37^ using a plasmid or PCR product and a synthetic ‘ligator” duplex oligonucleotide containing both dye and biotin modifications. Ligator was made by annealing two complementary oligonucleotides: 5′-CGATTAGGTCTCGGGCTAGTACTGGTTTCTAGAG/iCy5/GTTCCAAGCC/iBiodTCACGGCGGCCGCCCATCGAGACCGGTTAACC-3′ and 5′-GGTTA ACCGGTCTCGATGGGCGGCCGCCGTGAGGCTTGGAACCTCTAGAAACCAGTACTAGCCCGAGACCTAATCG-3′ (IDT).

For making template 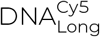, two identical Golden Gate Assembly reactions were carried out by mixing 7 *μ*l of each ∼ 20 *n*M DT4 plasmid and ∼ 20 *n*M ligator fragment with 1 *μ*l of Golden Gate Mix (New England Biolabs) in T4 DNA Ligase Buffer (New England Biolabs), in total volumes of 20 *μ*l. The mixtures were incubated for alternating cycles of 5 min at 37^*o*^C and 10 min at 16^*o*^C 35 times, followed by 5 min at 55^*o*^C. After the reaction, the ligase was inactivated for 10 min at 65^*o*^C. The resulting 40 *μ*l of reaction product was mixed with 4 *μ*l of T5 Exonuclease (New England Biolabs) in NEB Buffer 4, to a total of 50 *μ*l and incubated at 37^*o*^C for 30 min. The digestion was stopped by adding 15 *m*M of EDTA, and a Qiagen PCR Cleanup Kit was used to remove the cleaved nucleotides and enzymes.

For making template 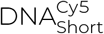, a linear DNA fragment was first amplified by PCR from plasmid pCDW114 (^16^, Addgene #70061), using primers 5′-GAAGGTCTCCAGCCGTACCAACCAGCGGCTTATC-3′ and 5′-CCGGG TCTCACCATACCCGCTGTCTGAGATTACG-3′. A Golden Gate Assembly reaction was carried out by mixing 2 *μ*l of 343 *n*M PCR product, 0.6 *μ*l of 1 *μ*M ligator and 1 *μ*l of Golden Gate Mix in T4 DNA Ligase Buffer, in a total volume of 20 *μ*l. The mixture was incubated for alternating cycles of 5 min at 37^*o*^C and 10 min at 16^*o*^C 35 times, followed by 5 min 55^*o*^C. After the reaction, the ligase was inactivated for 10 min at 65^*o*^C. The resulting reaction product was mixed with 1 *μ*l Exonuclease V (New England Biolabs) and 3 *μ*l of 10 *m*M ATP in NEB Buffer 4, in a total volume of 30 *μ*l, and incubated at 37^*o*^C for 30 min. Exonuclease V was then inactivated for 30 min at 70^*o*^C. Finally, a Qiagen PCR Cleanup Kit was used to remove the cleaved nucleotides and enzymes.

### Proteins

#### Fluorescent labeling of core RNAP

*E. coli* core RNAP with a SNAP tag on the C-terminus of *β*^*′* 38^ (RNAP-SNAP, gift from the Robert Landick lab) was labeled with SNAP-Surface 549, yielding RNAP^549^, as follows: 13.65 *μ*M SNAP-RNAP (core) and 45.5 *μ*M of SNAP-Surface 549 were mixed in a buffer containing 9 *m*M Tris-Cl^-^ pH 7.9, 5 *m*M MgCl_2_, 1 *m*M DTT, 20% glycerol, and 90 *m*M NaCl, and incubated for 30 min at room temperature. The sample was then mixed with an equivalent amount of Dilution buffer (11 *m*M Tris-Cl^-^ pH 8.0, 30% glycerol, 110 *m*M NaCl, 1 *m*M DTT), flash-frozen in liquid nitrogen and stored at -80^*o*^C.

#### Expression and purification of His-tagged *σ*^70^

His_6_-tagged *E. coli σ*^70^ (*σ*^70^) was overexpressed in T7 Express cells (New England Biolabs) as inclusion bodies from the pET-28a-*σ*^70^ overexpression plasmid^39^ by growing the cells at 37^*o*^C to an OD_600_ of ∼ 0.8, and then inducing by addition of IPTG to 0.4 *m*M. The temperature was decreased to 20^*o*^C and cells were left shaking at 200 rpm overnight. Cells were then harvested by centrifugation at 4^*o*^C, followed by sonication in Lysis buffer (50 *m*M Tris-Cl^-^, pH 7.9, 5 *m*M imidazole, 5% [v/v] glycerol, 233 *m*M NaCl, 2 *m*M EDTA, 10 *m*M *β*-mercaptoethanol) plus 1× cOmplete™protease inhibitor cocktail (Roche). The lysate was centrifuged at 22, 000 × *g* for 30 min at 4^*o*^C and the supernatant was discarded. To remove *E. coli* membrane and cell wall material, the pellet was resuspended in 10 *m*l of 2 M Urea Cleaning buffer (20 *m*M Tris-Cl^-^ pH 8.0, 500 *m*M NaCl, 2 M urea, 2% Triton X-100, 10 *m*M *β*-mercaptoethanol) and sonicated. The resulting sample was centrifuged again at 22, 000 × *g* for 30 min at 4^*o*^C and the supernatant was discarded. Four consecutive resuspension-centrifugation cycles were carried out, two of them in 2 M Urea Cleaning buffer, and the other two in Wash buffer (20 *m*M Tris-Cl^-^ pH 8.0, 500 *m*M NaCl, 7% glycerol, 20 *m*M imidazole, 10 *m*M *β*-mercaptoethanol) to remove Triton X-100 from the pellet. To solubilize and denature the protein, the washed pellet was resuspended in 6 M Guanidine Binding buffer (20 *m*M Tris-Cl^-^ pH 8.0, 500 *m*M NaCl, 5 *m*M imidazole, 6 M guanidine hydrochloride, 2 *m*M *β*-mercaptoethanol), stirred for 1 hour, and centrifuged at 22, 000 × *g* for 30 min at 4^*o*^C. The supernatant was passed through a 0.22 *μ*m filter and injected into a 1 *m*l HisTrap column (Cytiva Life Sciences), followed by a wash with Wash buffer supplemented with 6 M urea. Refolding of the bound protein was performed using a linear 1-hour-long 6 M to 0 M urea gradient in Wash buffer. The refolded protein was eluted with 500 *m*M imidazole in Wash buffer. The purified protein was dialyzed overnight into *σ*^70^ storage buffer (10 *m*M Tris-Cl^-^, pH 8.0, 30% [v/v] glycerol, 0.1 *m*M EDTA, 100 *m*M NaCl, 20 *μ*M ZnCl_2_, 1 *m*M MgCl_2_, 0.1 *m*M DTT) and aliquots were flash-frozen in liquid N_2_ and stored at -80^*o*^*C*.

#### Fluorescent labeling of *σ*^70^

An N-terminal His_6_-tagged single-cysteine derivative of *E. coli σ*^70^ (C132S C291S C295S S366C)^40^ (see Appendix S4) was overexpressed and purified following the same protocol used for *σ*^70^. The purified protein was concentrated 5× using an Amicon Ultra-0.5ml 30K filter by centrifuging for 11 minutes at 14, 000 × *g* at 4^*o*^C. For fluorescent labeling, the concentrated protein was mixed with Cy5-maleimide dye (Cytiva) (1:15 protein:dye ratio), incubated first for 10 min at room temperature, and then left overnight at 4^*o*^ C. The excess dye was then removed using a Centrispin 20 column (Princeton Separations). After addition of glycerol and BSA to 30% and 1 *m*g/*m*l respectively, the samples were flash-frozen in liquid N_2_, and aliquots were stored at -80^*o*^C.

#### Reconstitution of doubly-labeled holoenzyme

Cy5-*σ*^70^RNAP^549^ holoenzyme (see Appendix S4) was reconstituted by incubating 121 *n*M of RNAP^549^ and 280 *n*M of Cy5-*σ*^70^ for 30 min at 37^*o*^C.

### Colocalization Single-Molecule Spectroscopy (CoSMoS) Experiments

Single-molecule total internal reflection fluorescence microscopy was performed^41^ at excitation wave-lengths 532 and 633 *n*m, for observation of DNA^Cy5^ template (and/or Cy5-*σ*^70^) and RNAP^549^, respectively; focus was automatically maintained^42^. A stage heating device was used to keep the samples at 30^*o*^C. Single-molecule observations were performed in glass flow chambers (volume ∼ 30 *μ*l) passivated with a mPEG-SG2000:biotin-PEG-SVA5000 (Laysan Bio) 200 : 1 w/w mixture as described in^43^. Neutravidin (#21125; Life Technologies) was introduced at 220 *n*M in KO buffer (50 *m*M TrisOAc, 100 *m*M KOAc, 8 *m*M Mg(OAC)_2_, 27 *m*M NH_4_(OAc), 0.1 *m*g/*m*l bovine serum albumin (BSA) (#126615 EMB Chemicals), pH 8.0), incubated for 45 s, and washed out (this and all subsequent wash steps used two washes each of two chamber volumes of KO buffer). The chamber was then incubated with ∼1 *n*M Cy5-DNA (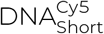 or 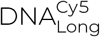) in KO buffer for ∼ 20 min and washed out with Imaging buffer (KO buffer supplemented with an O_2_ scavenging system: 4.5 *m*g/*m*l glucose, 40 units/*m*l glucose oxidase, and 1, 500 units/*m*l catalase^43^).

For the experiments to measure the dwell time of RNAP^549^ on DNA at different concentrations of *σ*^70^, ∼ 1 *n*M of RNAP^549^ was introduced into the chamber in Imaging buffer supplemented with 3.5% *w/v* PEG 8, 000 and 1 *m*g/*m*l BSA for ∼ 10 min. Image acquisition was performed by alternating 1 s exposures to 532 and 633 *n*m, at 450 and 200 *μ*W respectively (all laser powers were measured incident to the micromirror optics), and the flow chamber was washed with Imaging buffer supplemented with 3.5% *w/v* PEG 8, 000 and containing 0, 148 *n*M, 297 *n*M, 593 *n*M, 741 *n*M or 1.19 *μ*M of *σ*^70^. In the cases where the concentration of *σ*^70^ was lower than 1.19 *μ*M, the appropriate amount of *σ*^70^ storage buffer was added in replacement so that all experiments were performed at the same solute concentrations: 47.0 *m*M TrisOAc, pH 8.0, 93.0 *m*M KOAc, 7.4 *m*M Mg(OAc)_2_, 25 *m*M NH_4_(OAc), 3% *w/v* 8, 000 PEG, 0.2 *m*M Tris-Cl^-^, 2.0 *m*M NaCl, 0.4 *μ*M ZnCl_2_, 20 *μ*M MgCl_2_, 4.5 *m*g/*m*l glucose, 40 units/*m*l gluclose oxidase, 1, 500 units/*m*l catalase, 0.6% glycerol, 2 *μ*M EDTA, 0.1 *m*g/*m*l BSA, 2 *m*M DTT, 10 *n*M DTT-quenched Cy5.5 maleimide dye.

The experiments to measure dwell time of *σ*^70^RNAP on DNA (see Appendix S4) were performed similarly to the ones described in the previous paragraph, with three differences. First, a photobleaching step was performed after DNA surface attachment. Cy5 photobleaching was induced by 633 *n*m excitation at ∼ 1 mW in the presence of Imaging buffer without DTT. Second, instead of *σ*^70^, ∼ 1 *n*M of Cy5-*σ*^70^RNAP^549^ (which also contained an additional 1.5 *n*M Cy5-*σ*^70^) was introduced into the chamber in Imaging buffer supplemented with 3.5% *w/v* PEG 8, 000 and no subsequent wash was performed. The final composition of the solution was 47.5 *m*M of TrisOAc, pH 8.0, 95.1 *m*M KOAc, 7.4 *m*M Mg(OAc)_2_, 25.7 *m*M NH_4_(OAc), 3.3% *w/v* 8, 000 PEG, 0.1 *m*g/*m*l BSA, 1 *m*M DTT, 0.1 *m*M Tris-Cl^-^, 0.3% glycerol, 1 *m*M NaCl, 8 *μ*M MgCl_2_, 36 *n*M ZnCl_2_, 1.8 *μ*M EDTA, 1.1 *n*M SNAP-RNAP, 2.5 *n*M unreacted SNAP-Surface 549, 2.5 *n*M Cy5-*σ*^70^, 4.5 *m*g/*m*l glucose, 40 units/*m*l glucose oxidase, 1, 500 units/*m*l catalase. Third, image acquisition was performed by continuous exposure to 532 and 633 *n*m lasers, at 450 and 200 *μ*W respectively, at an acquisition rate of 1 frame per second.

### CoSMoS Data analysis

Analysis of CoSMoS video recordings was done using custom software and algorithms for mapping between wavelength channels, spatial drift correction, and detection of spot colocalization as described^41^. In each recording, we selected 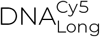 or 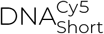 fluorescence spots that co-localized with RNAP^549^ spots at *t* = 0. For the selected DNA molecules, we computed RNAP^549^ fluorescence intensity time records by summing the intensity over 3 × 3 pixel squares centered at DNA molecule locations in each recorded frame. Fluorescence intensity values were corrected for background fluorescence and non-uniform illumination across the microscope field of view, yielding normalized values for spot intensities^44^. This allowed us to directly compare integrated intensity values for different spots located throughout the field of view. The numbers of decreasing intensity steps in the resulting time traces were counted to assess the initial number of RNAP^549^ molecules present at each DNA molecule location (Fig. S2A). Records that showed more than a single RNAP^549^ molecule bound at *t* = 0 were excluded from subsequent analysis. The times of the first image with no spot at each DNA location were taken to be the dwell times of the RNAP^549^ molecules present at the beginning of the recording. Spots that persisted until the end of the recording were separately counted as censored dwell times equal to the recording duration.

#### Fits to RNAP-DNA complex dwell time distributions

The probability distribution of RNAP-DNA complex dwell times measured in each individual experiment was modeled as the sum of two exponential terms

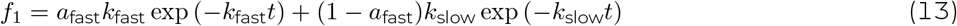

or the sum of three exponential terms

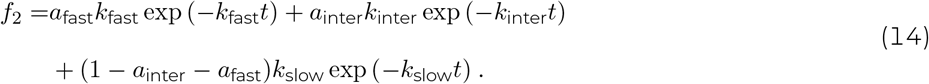

For each distribution, lifetimes of RNAP^549^ binding events that terminated by disappearance of the fluorescent spot, and those that were censored by the end of the experiment, were jointly fit using the maximum likelihood algorithm by an approach analogous to the one used in^45^. Confidence intervals were calculated by bootstrapping^41^.

#### Extraction of model parameters *k*_b_, *k*_off_, and *k*_off,s_

To get values for *k*_b_, *k*_off_, and *k*_off,s_, we globally fit dwell times collected at all concentrations of *σ*^70^ to the models in equation 13 (for [*σ*^70^] = 0) or equation 14 (for [*σ*^70^] *>* 0), where *k*_fast_ was explicitly constrained to depend on [*σ*^70^]:

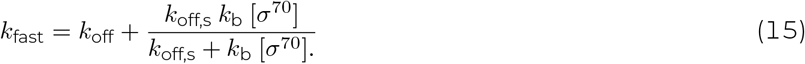

In this formulation, *k*_inter_ and *k*_slow_ are assumed to be independent of [*σ*^70^], and therefore were globally fit across distributions obtained for template 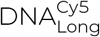. For the experiment performed with template 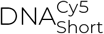, we fit independent values 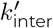 and 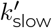 for these parameters. Censored data were treated using the same approach as described above for the individual experiment fits to dwell time distributions.

#### Fit to holoenzyme-DNA dwell time distribution

To account for non-specific binding of holoenzyme to the glass flow chamber (see Appendix S4), we first randomly selected locations on the chamber surface that did not contain DNA molecules. We then fit the distribution of dwell times for Cy5-*σ*^70^RNAP^549^ holoenzyme molecules bound at these non-DNA locations to a biexponential model

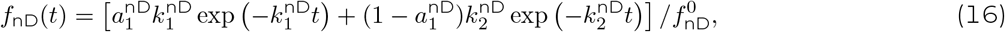

with

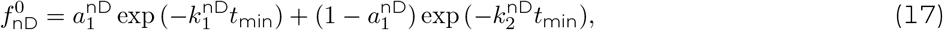

where *f*_nD_(*t*) is normalized so that it integrates to 1 over all dwell times *t* greater than the minimum detectable dwell time *t*_min_ = 0.2 s.

By analogy to^41^, the dwell time distribution for Cy5-*σ*^70^RNAP^549^ at DNA locations was then fit to the background-corrected model

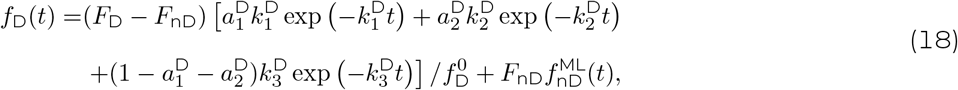

with

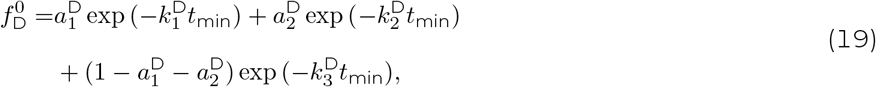

where *F*_D_ and *F*_nD_ represent the total binding frequency at DNA and non-DNA locations, respectively, and 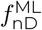 stands for *f*_nD_ evaluated on the maximum likelihood estimators obtained by fitting the non-DNA locations data. As before, we jointly fit the censored and uncensored data using the maximum likelihood method^45^.

### Simulation of search times

In order to calculate the mean search time ⟨*t*_f_⟩ required for the RNAP to find the secondary promoter, we used numerical simulations of the mechanism depicted in Fig. 1. In particular, we used the Gillespie algorithm^46^ to generate the stochastic trajectory of an RNAP molecule on a DNA molecule. In the simulation, the state of the system is characterized by the position of the RNAP on DNA and whether it is bound to *σ*^70^ or not. Initially, the RNAP is at position *x* = 0 (which corresponds to the position of the terminator from the primary transcription unit), and it is not bound to *σ*^70^. We then draw a time *t*_1_ at random from an exponential distribution with *p*(*t*) = *λ* exp (−*λt*) with *λ* = *k*_b_[*σ*^70^] + *k*_off_, and choose between two possible transitions: binding *σ*^70^ or dissociating from the DNA. Which of the transitions takes place is chosen at random according to their relative weights 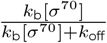 and 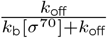. If the RNAP dissociates from the DNA, its attempt to find the secondary promoter is considered unsuccessful, and the simulation starts over with a new trial. If the RNAP binds *σ*^70^, the position on the DNA *x*_1_ at which binding occurs is dependent on the amount of diffusion away from the primary terminator. *x*_1_ is determined by drawing at random from a normal distribution with mean *μ* = 0 and variance var = 2*Dt*_1_. The time *t*_2_ that is required to diffuse from *x*_1_ to the secondary promoter located at *x*_*p*_ is then drawn at random from the first passage time density

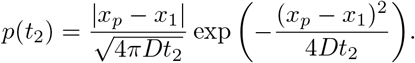

An RNAP-*σ*^70^ complex dissociation time *t*_3_ is drawn at random from an exponential distribution with *λ* = *k*_off,s_. If *t*_2_ < *t*_3_, the RNAP-*σ* complex is considered to have found the secondary promoter at time *t*_f_ = *t*_1_ + *t*_2_. If *t*_2_ *> t*_3_, the attempt to find the secondary promoter was unsuccessful. The whole process is repeated multiple times, generating a distribution of search times.

### Genome-wide analysis of terminator-promoter distances

Using the promoter and terminator annotations reported by Conway et al^29^, we measured the distance of each operon-ending terminator to the nearest operon initial promoter in the *E. coli* genome.

## ACKNOWLEDGMENTS

We thank Bob Landick, and Rachel Mooney, and for providing us with plasmids and proteins. We would like to thank members of the Landick, Kondev and Gelles labs for insightful discussion. We are grateful to Johnson Chung for help with microscopy, and with Liuyu Chen for help in plasmid preparation. This work was funded by grants from NIGMS (R01 GM081648 to J.G.), from NSF (DMR-1610737 to JK), and the Simons Foundation (to JK).

## AUTHOR CONTRIBUTIONS

D.T., L.F., J.G. and J.K. designed the research; D.T., K.I. and A.C. performed the research. D.T. analyzed the data; D.T., J.G., and J.K. drafted the manuscript and all authors contributed to writing the final version.

## Supporting Information for

## Appendix S1: Approximation of coupling distance 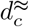 for the case [*σ*^70^] ≈ [*σ*^70^]_*c*_

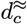 is defined in the main text as the distance for which the coupling efficiency *E* decays by a factor of 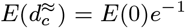, which in this case converts to

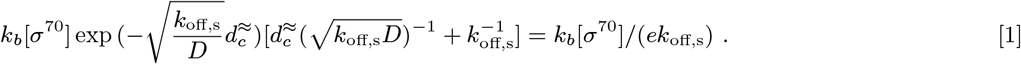

Defining 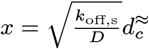, this reduces to the equation,

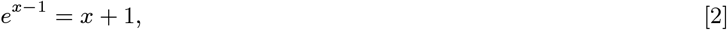

whose solution is given by

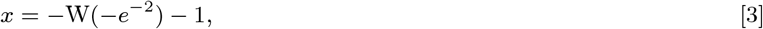

where W(*z*) is the product log function. The only real positive solution to eqn. 3 is given by

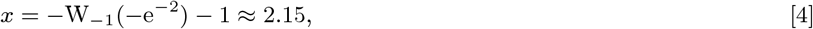

which gives 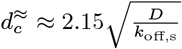.

## Appendix S2: General approximation of coupling distance

Values for model parameters *D, k*_b_[*σ*^70^], *k*_off_, and *k*_off,s_ were randomly drawn from log-uniform distributions in the ranges:

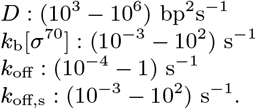

The characteristic coupling distance *d*_*c*_ (i.e., the distance for which the coupling efficiency decreases by a factor of *e*) was numerically calculated from Eq. 8, and plotted against the approximated value 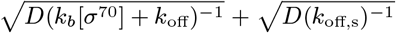 (Fig. S1), confirming that the coupling distance can be approximated by

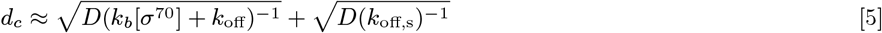

for all parameter values tested.

## Appendix S3: Selection of DNA templates that colocalized with a single RNAP^549^ molecule at *t* = 0

Because of the length of the DNA^Cy5^ templates employed (3,033 bp for 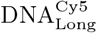 and 586 bp for 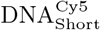), multiple RNAP^549^ could be bound to the same DNA molecule simultaneously. We restricted the analysis to those 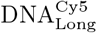 or 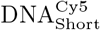 spots that colocalized with a single RNAP^549^ molecule at *t* = 0.

To quantify the number of RNAP^549^ molecules bound to each DNA, We used the integrated fluorescence time records. In the cases where the RNAP^549^ spots present at *t* = 0 disappeared during the duration of the experiment, the number of decreasing intensity steps was counted. Fig. S2A shows examples for one and two RNAP^549^ molecules present initially. DNA templates containing more than two RNAP^549^ molecules bound were rare.

For the cases in which the RNAP^549^ spots present at *t* = 0 did not disappear during the experiment, the absolute integrated intensity value was used to estimate the number of RNAP^549^ molecules present (Fig. S2B).

## Appendix S4: Experimental evidence for and interpretation of the intermediate and slow dwell time distribution components

In addition to the *σ*^70^-dependent component characterized by the dissociation rate *k*_fast_, we observed components in the RNAP-DNA dwell time distributions that did not change systematically with *σ*^70^ concentration. We investigated the origin of these *σ*^70^-independent components as follows:

Our preparation of RNAP^549^ showed low levels of *σ*^70^, indicating contamination of the core RNAP with *σ*^70^RNAP holoenzyme (Fig. S5). This raised the possibility that RNAP dwell time components with the *σ*^70^-independent rate constants *k*_inter_ and/or *k*_slow_ are due to *σ*^70^RNAP. To test this hypothesis,we measured the dwell times of *σ*^70^RNAP on DNA and compared them to dwell time components measured for the RNAP-*σ*^70^-DNA (RNAP-DNA complex that had bound *σ*^70^ from solution). Specifically, we performed a control experiment with doubly-labeled *σ*^70^RNAP holoenzyme (Cy5-*σ*^70^RNAP^549^) on template 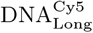 (Fig. S6A) and scored only dwell times at DNA locations where there was simultaneous colocalization of RNAP^549^ and Cy5-*σ*^70^ spots. This criterion excluded from our measurements dwell times of core RNAP molecules that did not have bound *σ*^70^. When the Cy5-*σ*^70^ fluorescent spot disappeared before the corresponding RNAP^549^ spot (e.g., Fig. S6B, right), the dwell time of the latter was measured.

In this experiment, roughly 10 − 15% of 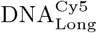 spots showed colocalization with more than one RNAP^549^ molecule bound simultaneously, often accompanied by Cy5-*σ*^70^, at some intervals during the recording. To simplify the interpretation of the data, we removed those records from the analysis, retaining only records that showed at most one core RNAP molecule at a time (e.g., Fig. S6B).

A fit of the resulting dwell time distribution (Fig. S6C) to a three-exponential background-corrected model (Eq. 18) (1) yielded the rate constants shown in Table S2. The fastest rate 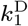 is approximately consistent with time constants previously observed for *σ*^70^ RNAP on nonspecific DNA (2). 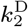 and 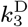 are approximately *k*_inter_ and *k*_slow_, respectively (Figs. S6D and E). This is consistent with the hypothesis that *k*_inter_ and *k*_slow_ are due to the minor *σ*^70^ contamination of the SNAP-RNAP sample used, and do not represent *σ*^70^-induced dissociation of RNAP from DNA.

The relative amount of the component with rate constant *k*_slow_, *a*_slow_ = 1 −*a*_fast_ −*a*_inter_, was roughly constant across experiments that used the same DNA preparation, even when [*σ*^70^] was different. Unexpectedly, *a*_slow_ was different between experiments using different preparations of the same DNA, and was almost absent in one of them (Fig. S7A). We propose that the events from this longest time component are due to stable binding of RNAP^549^ or *σ*^70^RNAP^549^ to some kind of imperfections in a fraction of the DNA molecules, such as nicks or gaps, which may be a consequence of the method used to prepare the DNAs and therefore will in general differ in abundance for different DNA preparations. In contrast, the fraction of the other (i.e., non-slow) dwells that are in the intermediate component, *a*_inter_*/*(*a*_inter_ + *a*_fast_), was roughly constant for two different 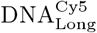 preparations and for the two different DNA sequences, 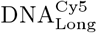 and 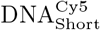 (Fig. S7B). A possible explanation for the binding events with characteristic dissociation rate *k*_inter_ is that they arise from holoenzyme binding to rare tight binding sequences (3–5) that may be present in both of the DNA circles tested. More experiments will be needed to test this hypothesis.

**Fig. S1.**
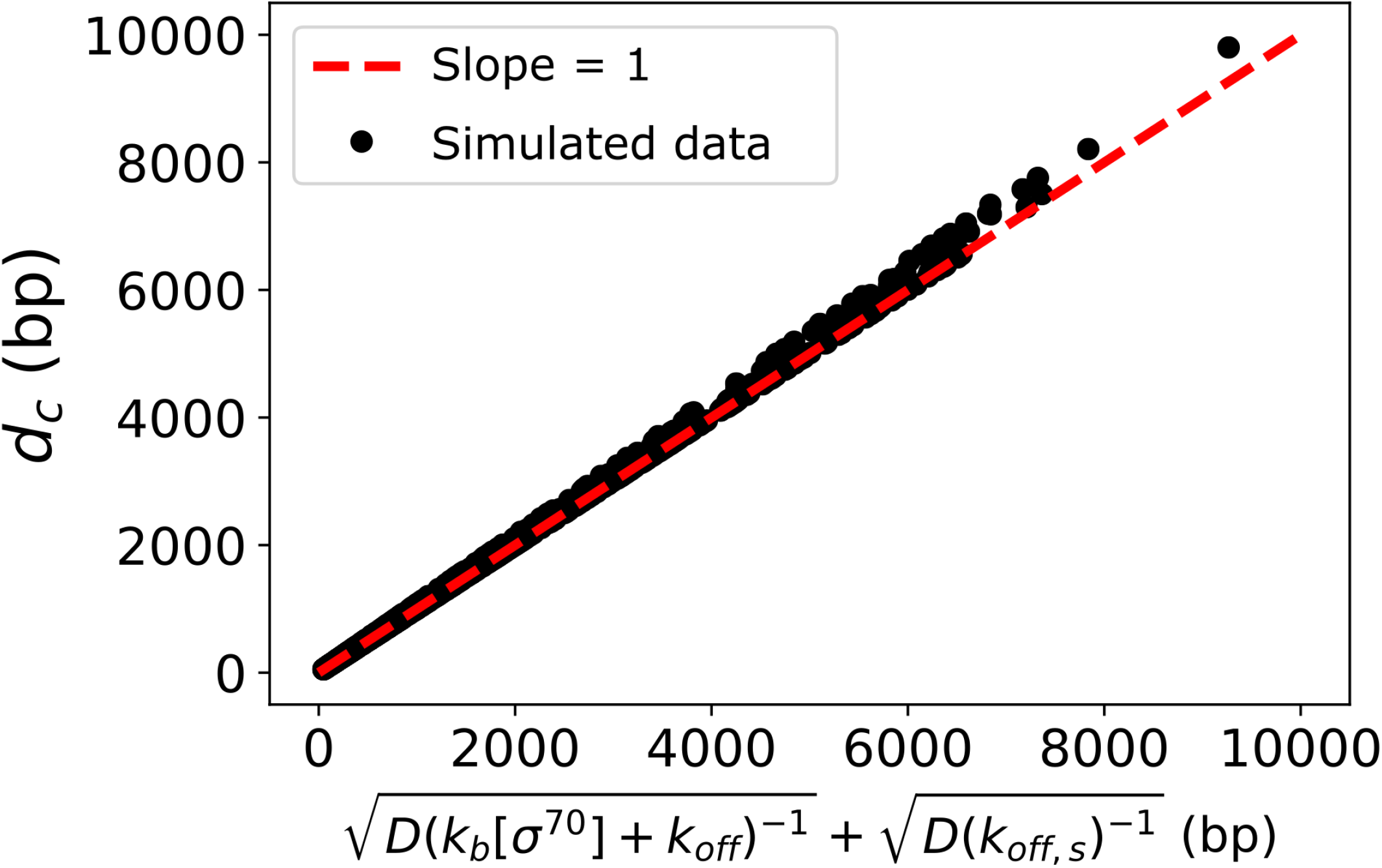
Approximation of coupling distance *d*_*c*_. Coupling distance was numerically calculated from Equation 8 as the value of *d* over which the efficiency *E* decreases by a factor of *e*, using parameter values chosen at random (black dots, see Appendix S2). That exact calculation agrees well with our approximate expression for *d*_*c*_ (red line), confirming that *d*_*c*_ can be approximated by 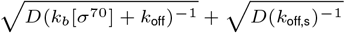.

**Fig. S2.**
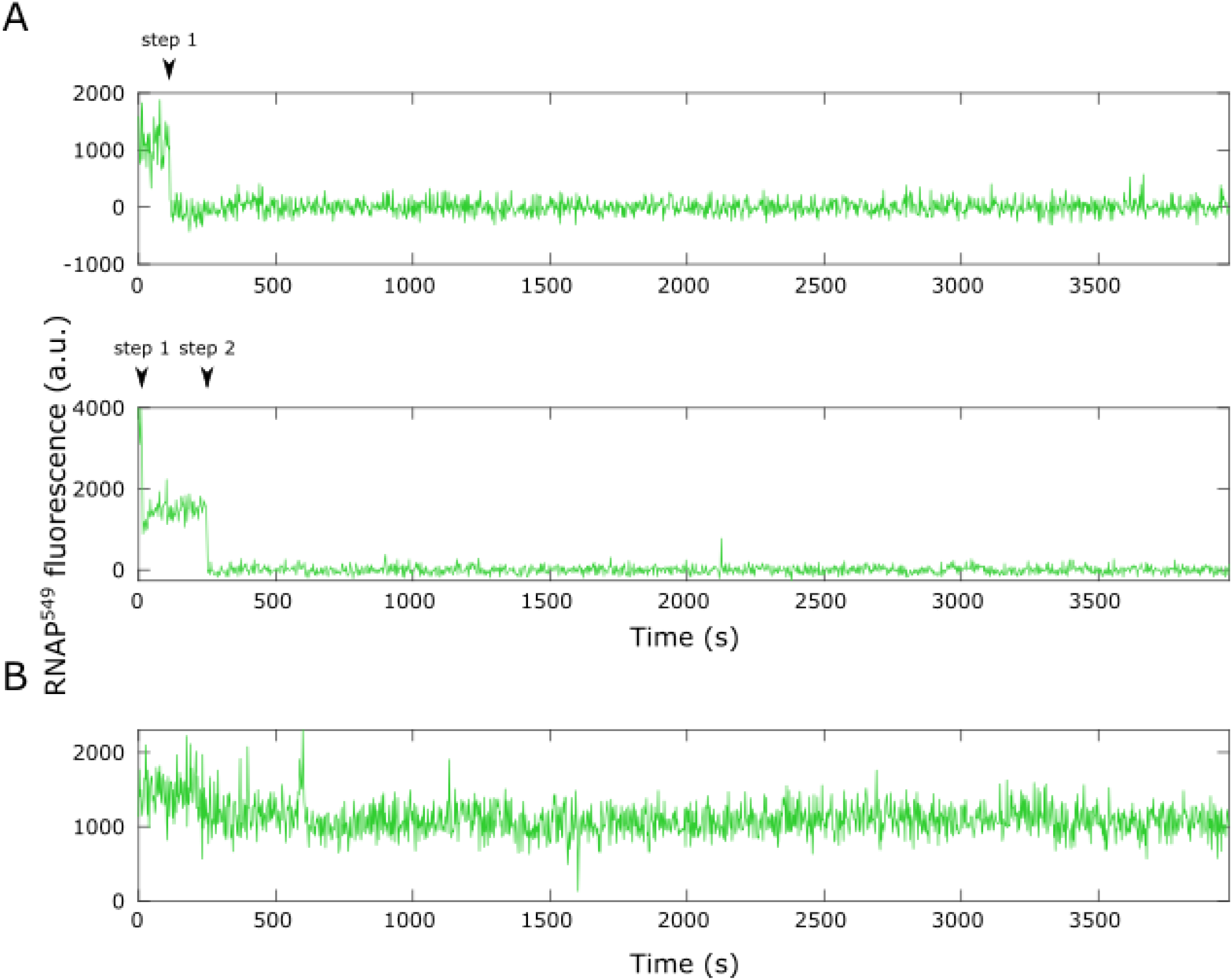
Counting the number of RNAP^549^ molecules bound to an individual DNA molecule at *t* = 0. **A**. Records showing one (top) and two (bottom) RNAP^549^ molecules that dissociate or photobleach during the experiment. **B**. Record showing a single RNAP^549^ molecule that lasts throughout the experiment, as indicated by the roughly constant normalized fluorecesce intensity of ∼1, 000. In all records, zero fluorescence corresponds to the diffuse background fluorescence detected in the absence of a DNA-colocalized RNAP^549^ molecule.

**Fig. S3.**
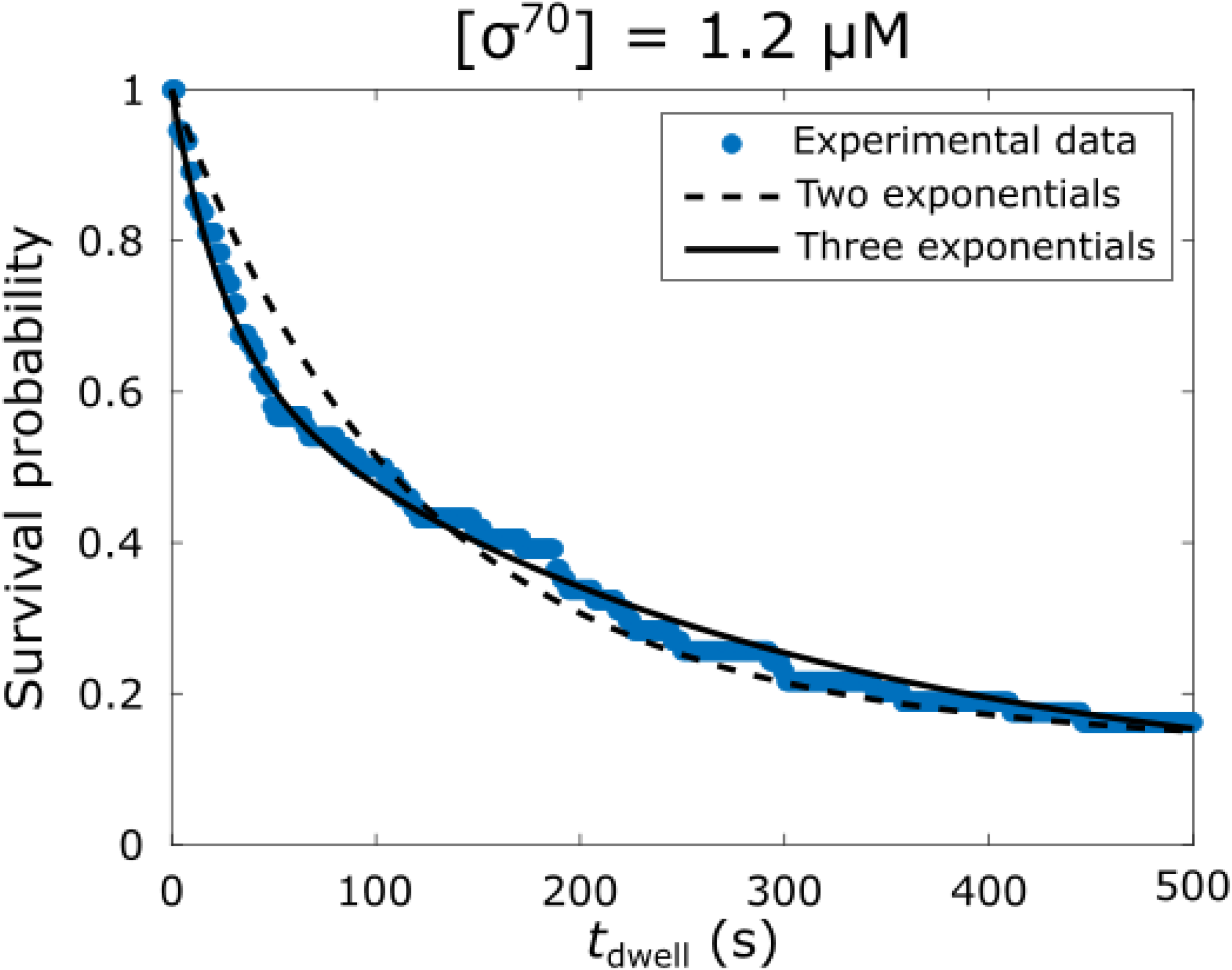
Comparison between two-exponential and three-exponential fits to an example RNAP^549^ −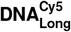 dwell time distribution. The distribution obtained at 1.19 *μ*M *σ*^70^ (blue) was fit to a sum of two exponentials model (Eq. 13 with *a*_fast_ = 8.2(6.9−9.2)*×*10^−1^, *k*_fast_ = 8.7(6.0−14.2)*×*10^−3^ s^−1^, *k*_slow_ = 4.7(1.2−10.0)*×*10^−4^ s^−1^; dashed line) and a sum of three exponentials model (Eq. 14 with *a*_fast_ = 3.4(2.4 − 4.9)*×* 10^−1^, *k*_fast_ = 3.9(2.4 − 5.4)*×* 10^−2^ s^−1^, *a*_inter_ = 5.8(4.4 − 6.7)*×* 10^−1^, *k*_inter_ = 3.9(2.5 − 5.2) *×* 10^−3^ s^−1^, *k*_slow_ = 1.6(0 − 3.5) *×* 10^−4^ s^−1^; solid line). The values are presented with 68% C.I.s.

**Fig. S4.**
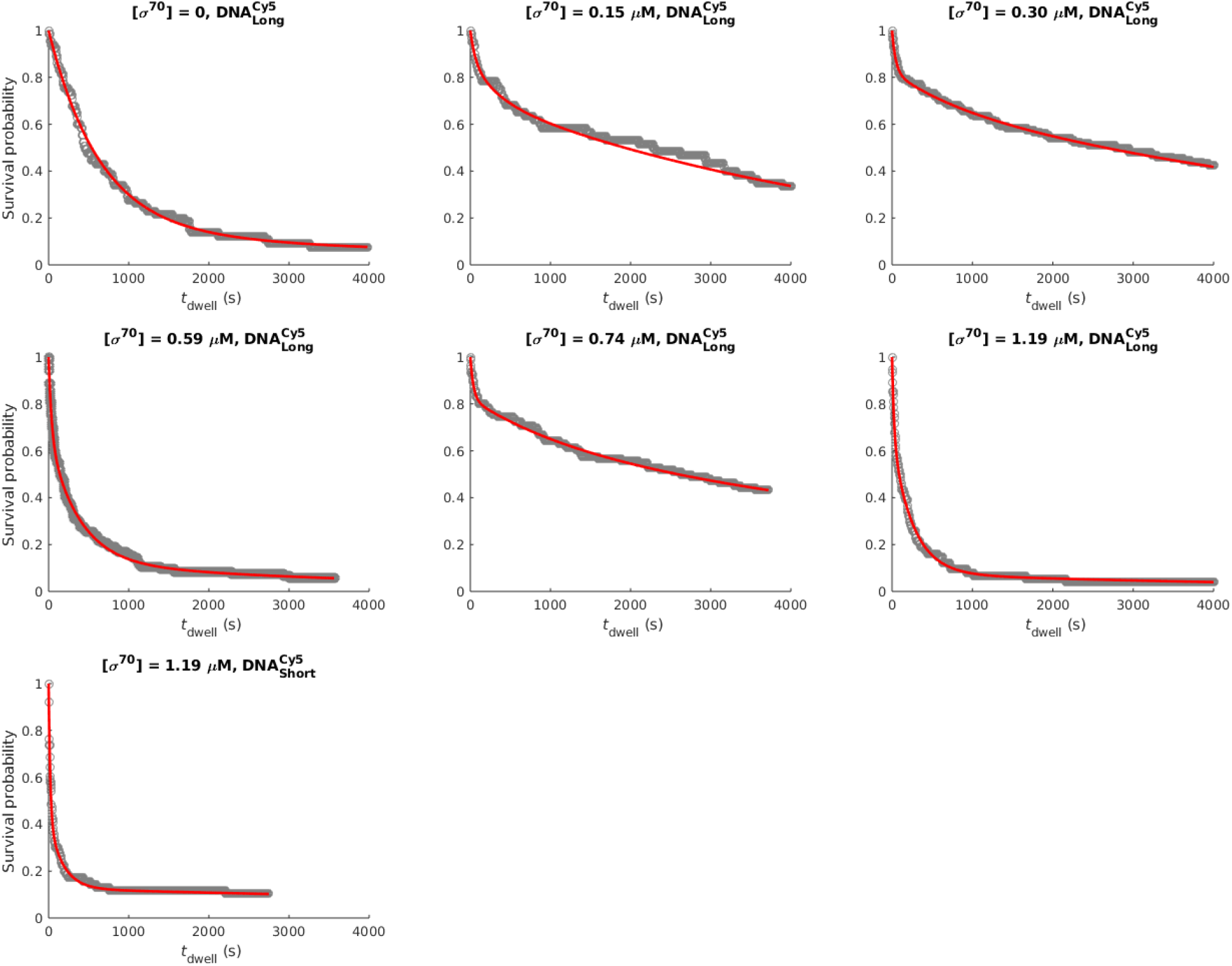
RNAP^549^ −DNA dwell time distributions and local fits with two different DNA constructs in the presence of different concentrations of *σ*^70^. Each distribution was fit to the sum of two exponentials (for [*σ*^70^] = 0) or the sum of three exponentials (for [*σ*^70^] *>* 0). The fit parameters are reported in Table 1.

**Fig. S5.**
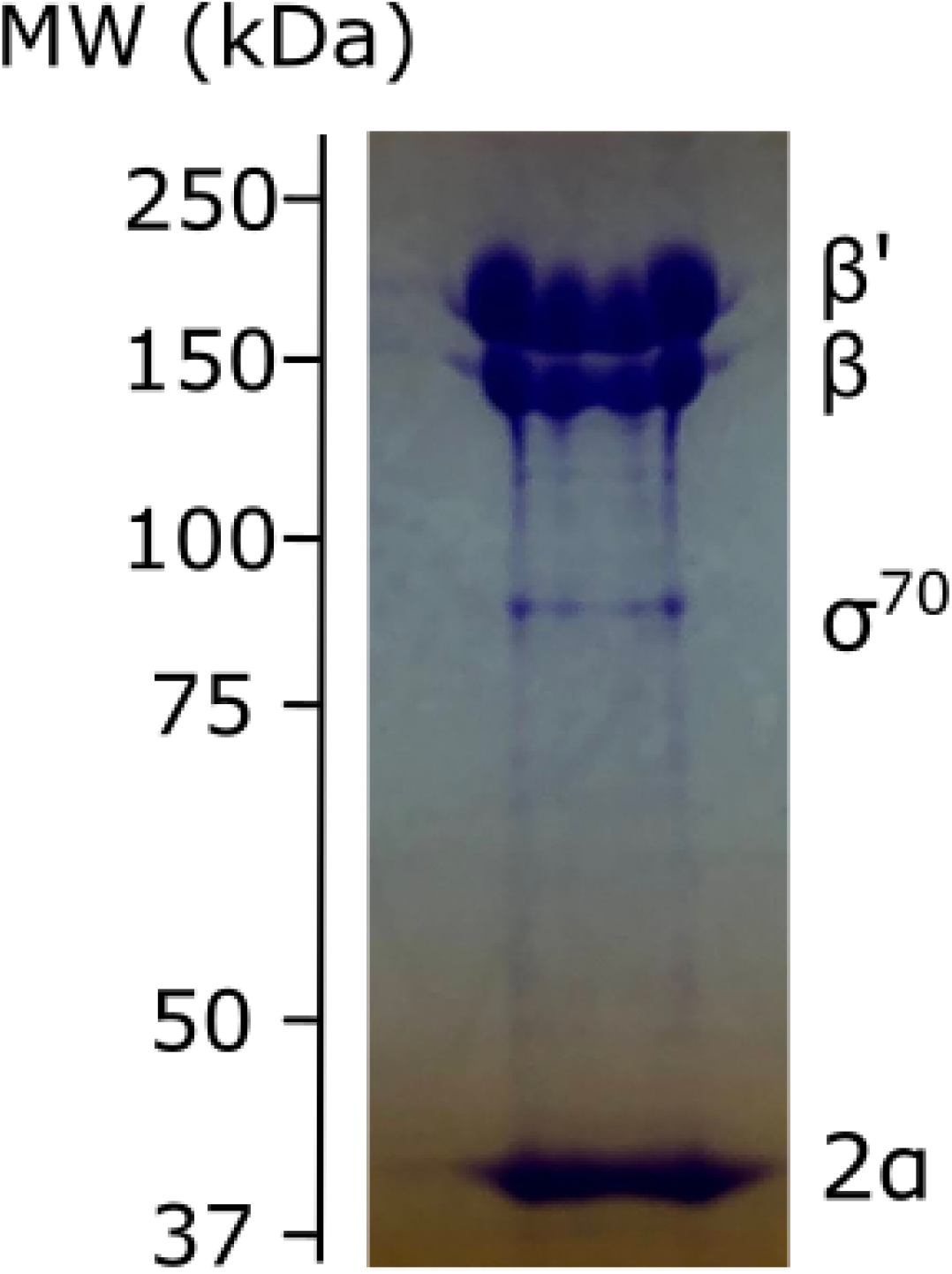
Extent of *σ*^70^ contamination in the RNAP-SNAP preparation used in this study. Densitometry of the Coomassie-Blue stained SDS-PAGE gel indicates the presence of *σ*^70^ at ∼ 7.5 mole percent of core.

**Fig. S6.**
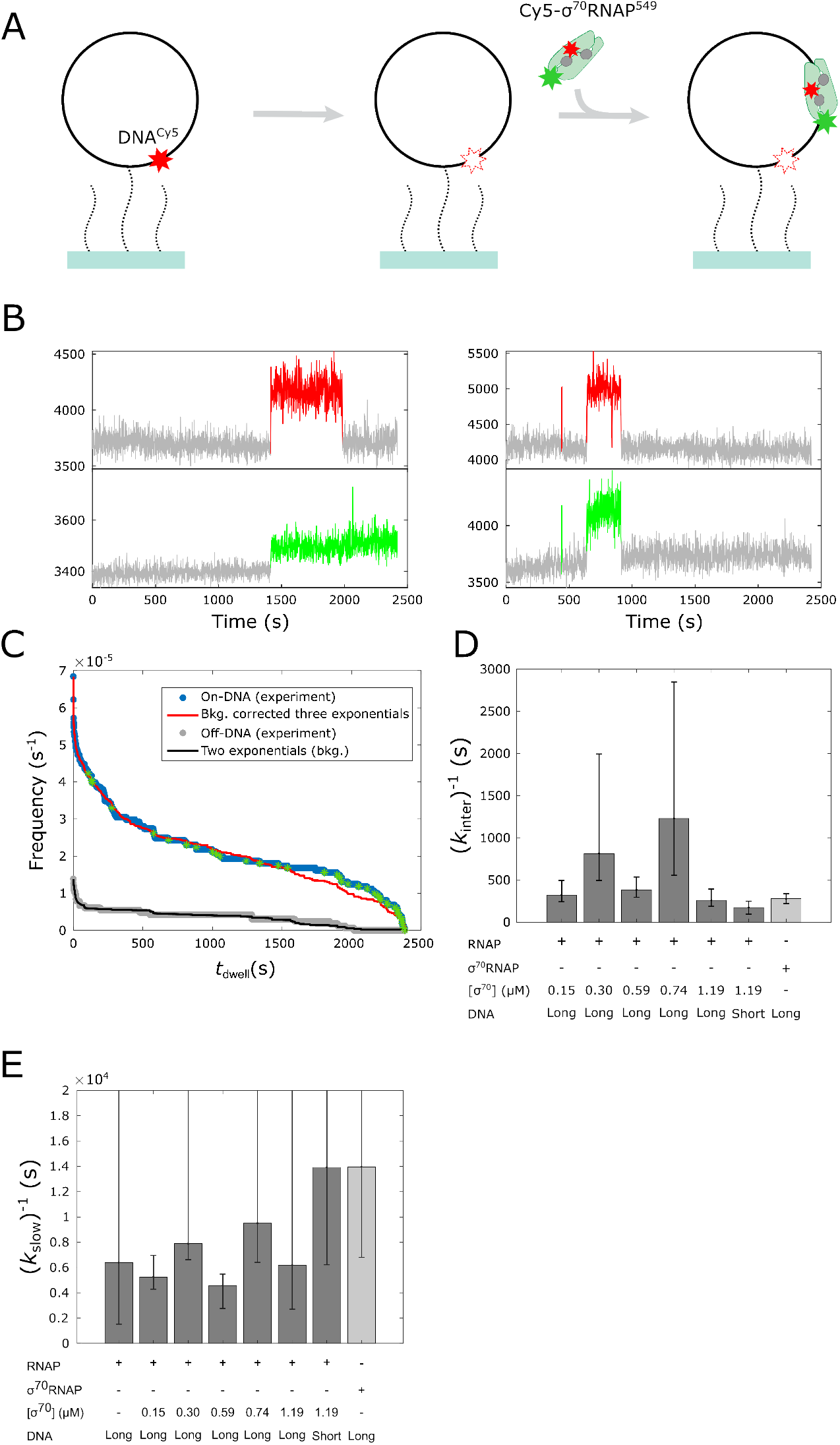
Control experiment with doubly-labeled *σ*^70^RNAP holoenzyme. See Appendix S4. **A**. Experiment design. Fluorescently labeled, promoterless circular DNA molecules (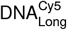; black circles with red stars) were tethered to the surface of a glass flow chamber (blue) through polyethylene glycol linkers (dotted black curves). After recording the position of the DNA molecules, the fluorescent dye was photobleached (white star). Doubly-labeled RNAP holoenzyme Cy5-*σ*^70^RNAP^549^ was added and two-color fluorescence time records were acquired. **B**. Examples of two-color Cy5-*σ*^70^RNAP^549^ fluorescence time records recorded at the locations of two individual DNA molecules. Time intervals in which the fluorescently labeled protein subunits were detected are indicated (color). **C**. Cumulative frequency distributions for Cy5-*σ*^70^RNAP^549^ molecules binding to 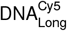 (blue) or binding to non-DNA positions (gray) with a dwell time greater than or equal to the specified value. Stars indicate censored dwell times that last until the end of the experiment. Figure also shows a two-exponential distribution function (black) fit to the non-DNA data and a three-exponential background-corrected distribution function (red) fit to the DNA data (1). Distribution function plots were numerically simulated to include the effects of censoring. The fit parameters are summarized in Table S2. **D**. The characteristic durations (with 68% C.I.s) of intermediate-length binding events from experiments with unlabeled *σ*^70^ (dark gray; Fig. 3A; *k*_inter_ in table 1) is consistent with the characteristic duration of intermediate-length specific binding events of Cy5-*σ*^70^RNAP^549^ holoenzyme (light gray; see panel C; *k*_2_ in Table S2). **E**. Same as D, but for long-duration time constants. While C.I. lower bounds are well determined, most of the C.I. upper bounds are poorly defined because the reciprocal rate constants exceed the durations of the experiments, which range from 2, 749 s to 4, 570 s.

**Fig. S7.**
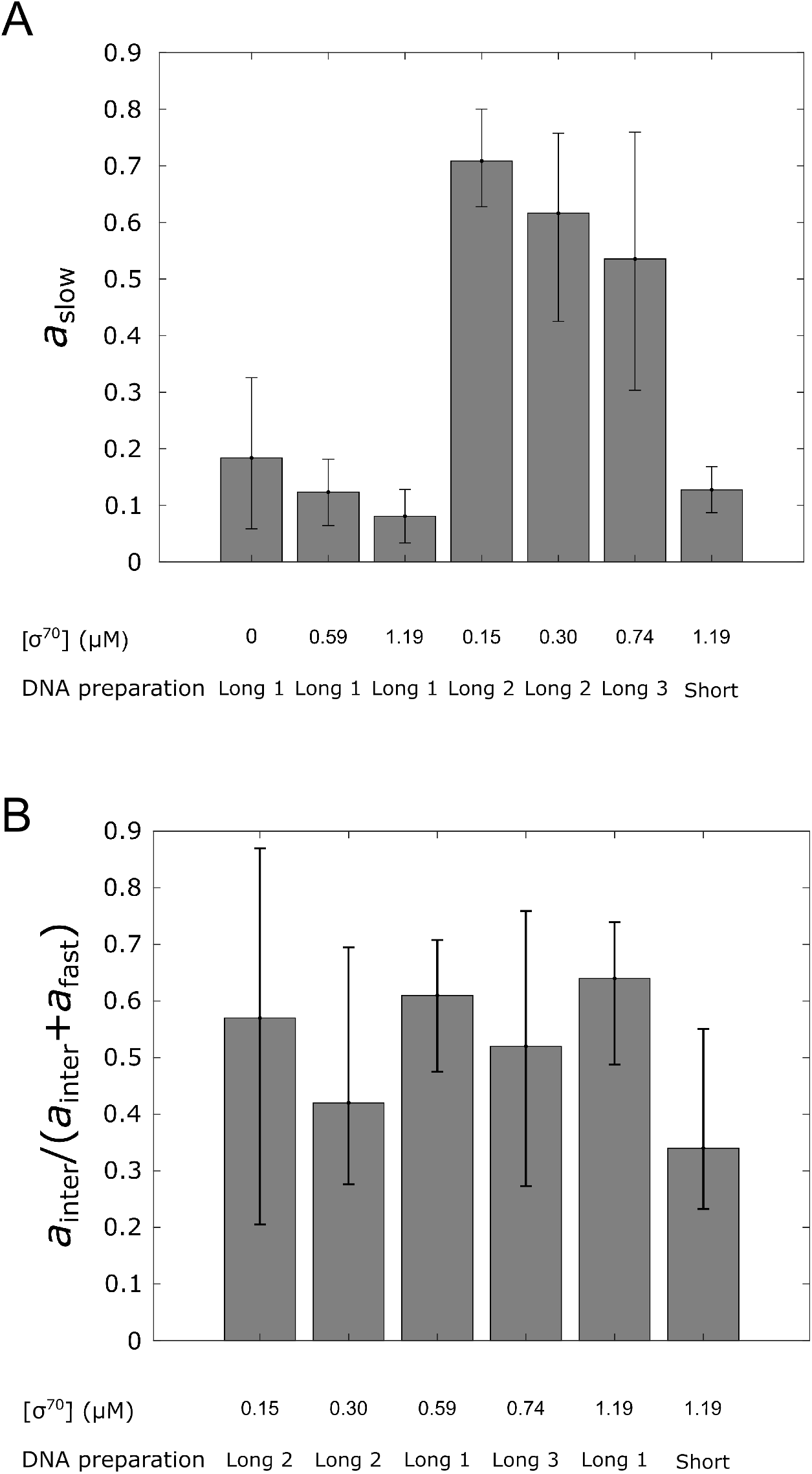
Amplitudes of the slow and intermediate dwell time components in experiments with different DNA preparations and different [*σ*^70^]. **A**. Relative amplitude of the slow dissociation component *a*_slow_ for experiments performed using three different preparations of circular DNA 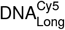, all built following the same protocol, and one preparation of 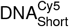. The value of *a*_slow_ = 1 − *a*_inter_ − *a*_fast_ (Eq. 14) depends on the DNA templates preparation used, supporting the hypothesis that the longest binding events are due to stable binding to imperfections in some of the DNA molecules. **B**. Fraction of the non-slow dwells that are in the intermediate component, as given by a_inter_*/*(a_inter_ + a_fast_) in the same experiments as in (A). This quantity is roughly constant across experiments on all DNA constructs and preparations.

**Fig. S8.**
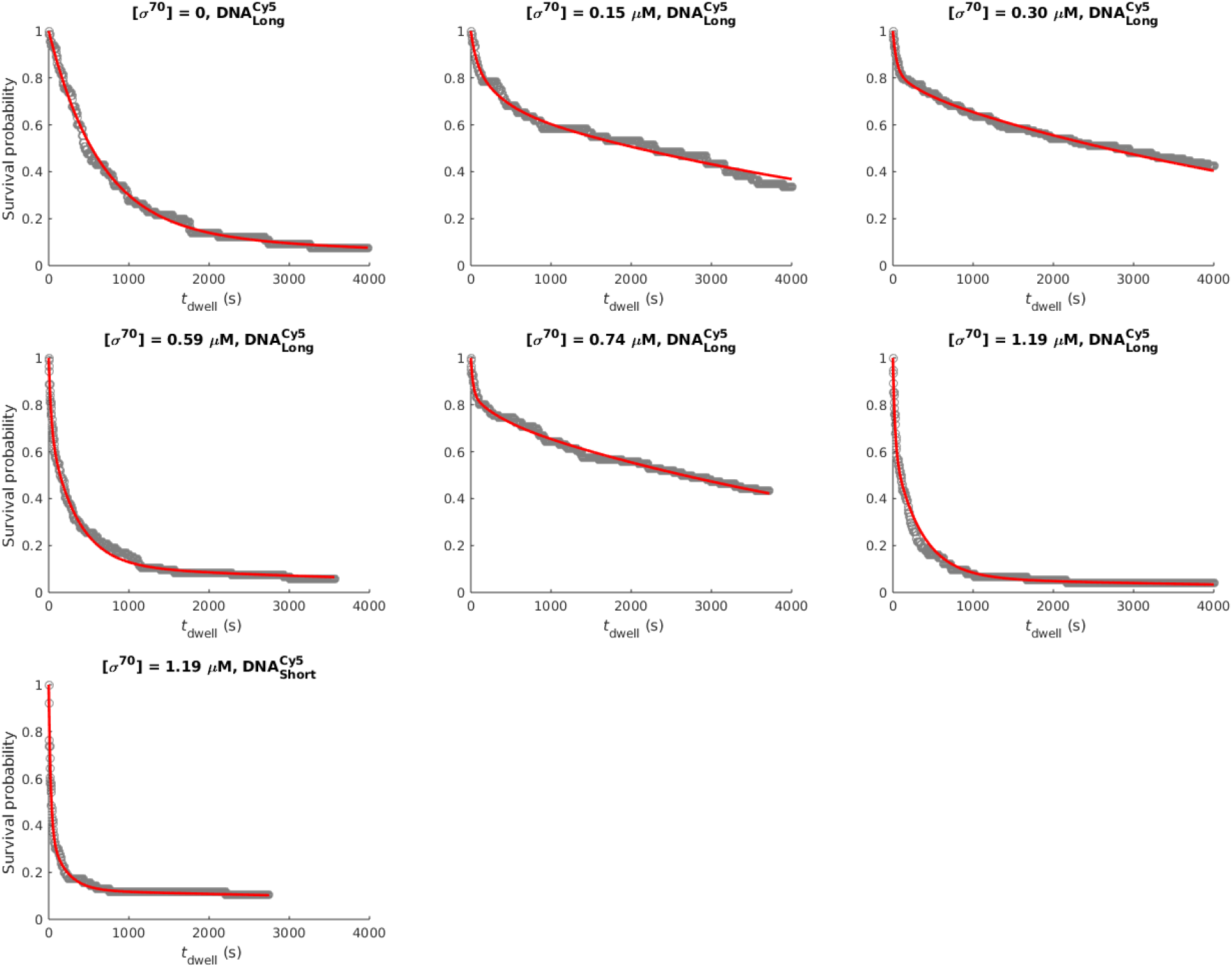
RNAP^549^ −DNA dwell time distributions and global fits with two different DNA constructs in the presence of different concentrations of *σ*^70^. Displayed fits (red) are to the global model in which *k*_inter_ and *k*_slow_ are fixed across experiments, and *k*_fast_ is constrained to a hyperbolic dependence on *σ*^70^ concentration. This contrasts with Fig. S4 in which we used independent local parameter values for each dataset. Parameters for the fits shown here are summarized in Table 2 and Table S3. Experimental data (gray) are the same as in Fig. S4.

**Table S1.** Fraction of DNA and non-DNA locations that had one RNAP^549^ molecule bound at *t* = 0.

| $[\sigma^{70}](\mu\text{M})$ | Fraction of locations | |
| --- | --- | --- |
|  | DNA | non-DNA |
| 0 | 18% | 1.8% |
| 0.15 | 15% | 0.5% |
| 0.30 | 31% | 0.9% |
| 0.59 | 17% | 1.6% |
| 0.74 | 15% | 0.7% |
| 1.19 | 25% | 1.3% |

The non-DNA binding was measured by observing the binding of RNAP<sup>549</sup> molecules at random positions in the microscope field of view that contained no visible DNA<sup>Cy5<sub>Long</sub></sup> molecules.

**Table S2.**
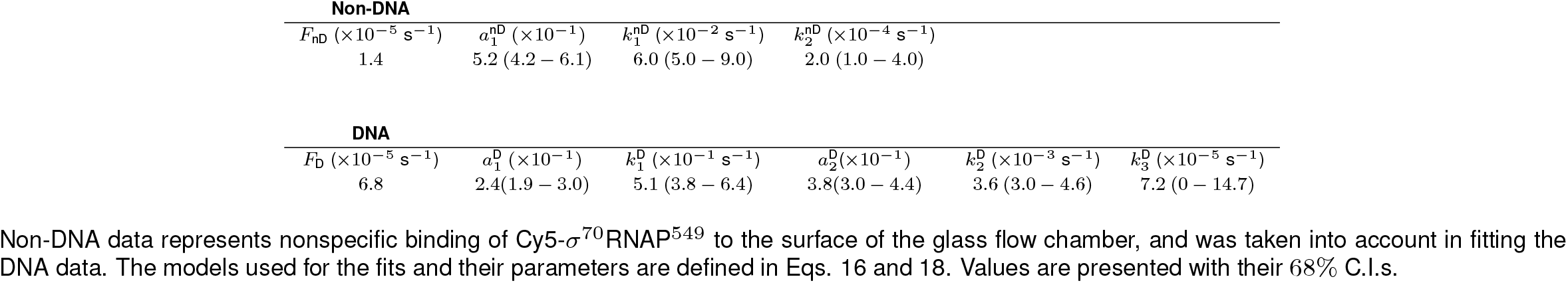
Fit parameters for distribution of Cy5-*σ*^70^RNAP^549^ dwell times on 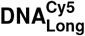.

**Table S3.**
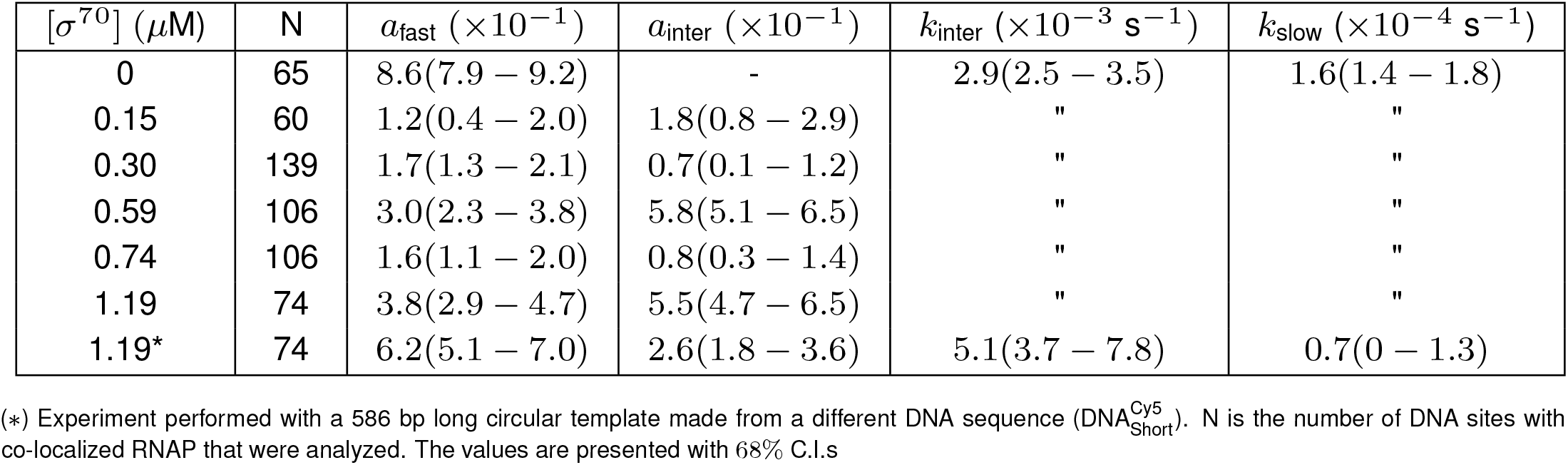
Fit parameters for survival lifetime probability distributions obtained by globally fixing *k*_inter_ and *k*_slow_ and considering *k*_fast_ dependence on [*σ*^70^].

